# A phylogenetic approach to explore the *Aspergillus fumigatus* conidial surface-associated proteome and its role in pathogenesis

**DOI:** 10.1101/2023.08.22.553365

**Authors:** Clara Valero, Camila Figueiredo Pinzan, Patrícia Alves de Castro, Norman van Rhijn, Kayleigh Earle, Hong Liu, Maria Augusta Crivelente Horta, Olaf Kniemeyer, Thomas Krüger, Annica Pschibul, Derya Nur Coemert, Thorsten Heinekamp, Axel A. Brakhage, Jacob L. Steenwyk, Matthew E. Mead, Antonis Rokas, Scott G. Filler, Nathalia Gonsales da Rosa-Garzon, Hamilton Cabral, Endrews Deljabe, Michael J. Bromley, Claudia B. Angeli, Giuseppe Palmisano, Ashraf S Ibrahim, Sara Gago, Thaila F. dos Reis, Gustavo H. Goldman

## Abstract

*Aspergillus fumigatus*, an important pulmonary fungal pathogen causing several diseases collectively called aspergillosis, relies on asexual spores or conidia for initiating host infection. Here, we used a phylogenomic approach to compare proteins in the conidial surface of *A. fumigatus*, two closely related non-pathogenic species, *Aspergillus fischeri* and *Aspergillus oerlinghausenensis*, and the cryptic pathogen *Aspergillus lentulus*. After identifying 62 proteins uniquely expressed on the *A. fumigatus* conidial surface, we deleted 42 genes encoding conidial proteins. We found deletion of 33 of these genes altered susceptibility to macrophage killing, penetration and damage to epithelial cells, and cytokine production. Notably, a gene that encodes glycosylasparaginase, which modulates levels of the host pro-inflammatory cytokine IL-1β, is important for infection in an immunocompetent murine model of fungal disease. These results suggest that *A. fumigatus* conidial surface proteins and effectors are important for evasion and modulation of the immune response at the onset of fungal infection.

## Introduction

Pulmonary fungal diseases caused by the environmental mold *Aspergillus fumigatus* lead to significant morbidity and mortality. In patients with weakened lung defenses arising from immunosuppression, a chronic respiratory condition, or a prior respiratory infection, asexual spores or conidia of *A. fumigatus* can evade the lung defenses, germinate, and cause disease ^1^. However, our understanding of what makes *A. fumigatus* a successful pathogen compared to other species remains incomplete. Current evidence suggests that several different *Aspergillus* species, including *A. fumigatus*, likely independently evolved the ability to cause human disease ^2^ (**Figure 1A**), raising the hypothesis that species-specific genes contribute to disease.

**Figure 1.**
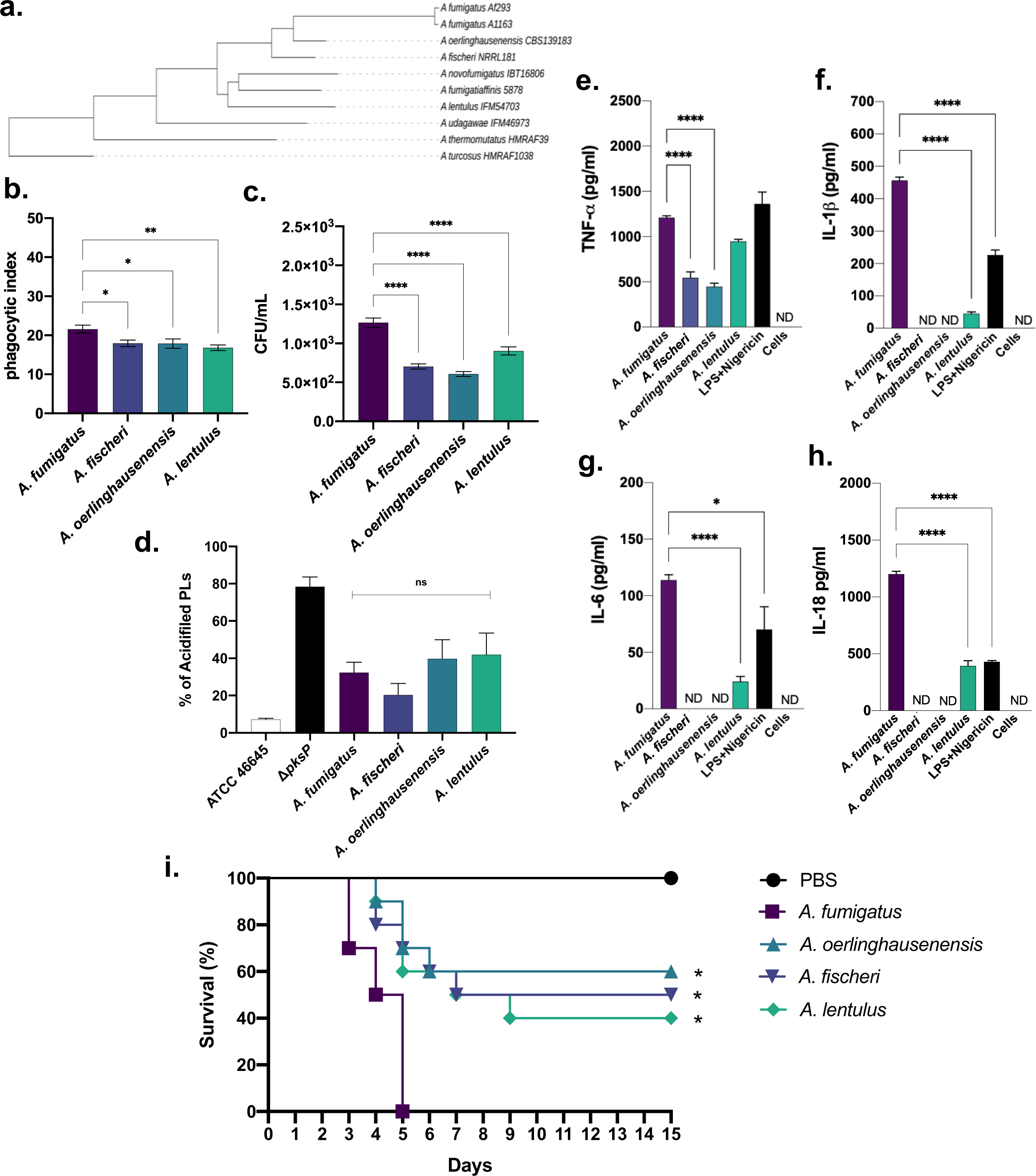
*A. fumigatus* has increased virulence and cytokine production by BMDMs than the other three species. **a.** Phylogeny of *Aspergillus* section *Fumigati* representative species constructed from concatenation analysis of a 5,215-gene data matrix. Branch lengths correspond to nucleotide substitutions / site. Adapted from ^58^. **b and c.** Phagocytic index and viability of conidida after exposure to BMDMs. **d.** Percentage of acidifed phagolysosomes (PLs). **e to h.** Cytokine production elicited by Afu, Afi, Aoer, and Ale conidia by BMDMs. **i.** Survival curves for *A. fumigatus* (Afu), *A. fischeri* (Afi), *A. oerlinghausenensis* (Aoer), and *A. lentulus* (Ale) in a chemotherapeutic murine model of Invasive Pulmonary Aspergillosis (IPA).

The early stages of disease—marked by the interaction between the inhaled conidia and the host—may prove insightful for unraveling *A. fumigatus* pathogenicity. Among studies of other microbial pathogens, there has been an increasing focus on conidial surface protein characterization since they mediate the first encounter with the host immune system. The conidial cell wall comprises a β-1,3-glucan and chitin body, which is covered by rodlet and melanin layers, where proteins are anchored ^3, 4^. These and other surface proteins play pivotal roles in morphogenesis, resistance to environmental stressors, substrate adherence, and virulence ^5^. The hydrophobin RodA is a significant component of the rodlet layer and is essential for cell wall physical resistance and permeability, preventing immune recognition ^6, 7^. Another conidial surface protein, CcpA, is essential to maintain the correct surface structure and prevent immune recognition ^8^. Recently, HscA has been demonstrated to anchor human p11 on phagosomal membranes, rewiring the vesicular trafficking to the non-degradative pathway, allowing the escape of conidia ^9^. Other studies aimed to identify potential vaccine candidates, allergens, and biomarkers for diagnosis or characterization of *A. fumigatus* cell wall dynamics under different stresses or biological processes^10–12^.

Despite the importance of the conidia for disease, a comprehensive examination of conidial surface proteins and their contribution to pathogenicity in *A. fumigatus* and closely related species is lacking. Here, we analyzed the conidial surface proteome of *A. fumigatus* and three closely related species: (i) *Aspergillus lentulus*, which is a cryptic pathogen and close relative of *A. fumigatus* ^13^; (ii) *Aspergillus oerlinghausenensis,* the closest known relative of *A. fumigatus,* which is azole-resistant but non-pathogenic ^14, 15^; and (iii) *Aspergillus fischeri*, a close relative of *A. fumigatus* that is less virulent in several animal models and rarely causes disease in humans ^16^ (**Figure 1a**).

## Results

***A. fumigatus* has increased virulence in a chemotherapeutic murine model of Invasive Pulmonary Aspergillosis (IPA) and elicits higher cytokine production by bone marrow-derived macrophages (BMDMs)**. To understand whether the four species have *in vivo* and *in vitro* differences in pathogenicity, we comparatively analyzed the capacity of *A. fumigatus*, *A. fischeri* (Afi), *A. oerlinghausensis* (Aoe), and *A. lentulus* (Ale) to infect macrophages and susceptible hosts. *A. fumigatus* conidia are less engulfed and killed by Bone Marrow derived macrophages (BMDMs) than Afi, Aoe, and Ale (**Figures 1b and 1c**). *A. fumigatus* conidia are recognized by macrophages and are intracellularly degraded by the endocytic pathway through the fusion of conidia-containing phagosomes and lysosomes, forming an acidic phagolysosome (PL) ^17^. Melanin can facilitate evading this mechanism, as the *A. fumigatus* Δ*pksP* mutant (which lacks a key enzyme involved in melanin biosynthesis) induces higher levels of PL acidification than the corresponding wild-type strain ATCC46645 (**Figure 1d**). However, there are no differences in PL acidification in *A. fumigatus*, Afi, Aoer, and Ale, indicating that these four species’ mechanisms of inhibiting PL acidification are conserved (**Figure 1d**). In contrast, *A. fumigatus* conidia induced higher levels of four cytokines—TNF-α, IL-6, IL-1β, and 1L-18—in BMDMs than the other three species (**Figures 1e to 1h**). Interestingly, *A. fumigatus* and Ale uniquely induced production of inflammasome cytokines IL-1β and IL-18, while Afi and Aoer conidia led to the production of lower levels of TNF-α than *A. fumigatus* and Afi conidia (**Figures 1e to 1h**).

Virulence of each species was assessed in a clinically relevant chemotherapeutic BALB/c murine model of IPA. The *A. fumigatus* strain was significantly more virulent than Afi, Aoe, and Ale strains (*p*-value > 0.005, **Figure 1i**) and killed all mice after 5 days post-infection (d.p.i.). However, while there was no statistical difference among the other species, an increased virulence caused by Ale strain was observed when compared to the Aoe and Afi strains (40% vs. 50-60% of survival at 15 d.p.i., **Figure 1i**).

Taken together, these results strongly indicate that *A. fumigatus* is more virulent in a chemotherapeutic murine model of IPA, is more efficiently recognized by BMDMs, and can elicit higher levels of cytokine production than the other three species.

### The conidial surfome of *A. fumigatus* contains 62 unique proteins

Variation in host recognition observed between *A. fumigatus* and the other three species raises the hypothesis that conidial surface proteins may underlie differences in host recognition variation. To test this hypothesis, we conducted a trypsin-shaving proteomic analysis and identified *A. fumigatus* unique proteins on the conidial surface. To do so, samples of resting (0h) and swollen (4h, 37°C) conidia from the four *Aspergillus* species included in the study (*A. fumigatus*, Afi, Aoe, and Ale; **Figure 2a**) were collected and processed for LC-MS/MS analysis as described in ^5^ (**Figure 2b**). A total of 354 proteins were identified in *A. fumigatus* conidial surface [193 in resting and 161 in swollen conidia, and 122 (52.6%) shared between the two conditions]. Similarly, in Afi, 594 proteins were identified as part of the conidial surfome [309 in resting and 285 in swollen conidia, and 204 (52.3%) shared between the two conditions]. For Ale, we identified 1,027 conidial surface proteins [285 in resting and 742 in swollen conidia, but only 240 (30.1%) shared between the two conditions]. Finally, 763 proteins were identified in Aoe conidial surface [393 in resting and 370 in swollen conidia, and 269 (54.5%) shared between the two conditions] (**Supplementary Figures S1; Supplementary Table S1**).

**Figure 2.**
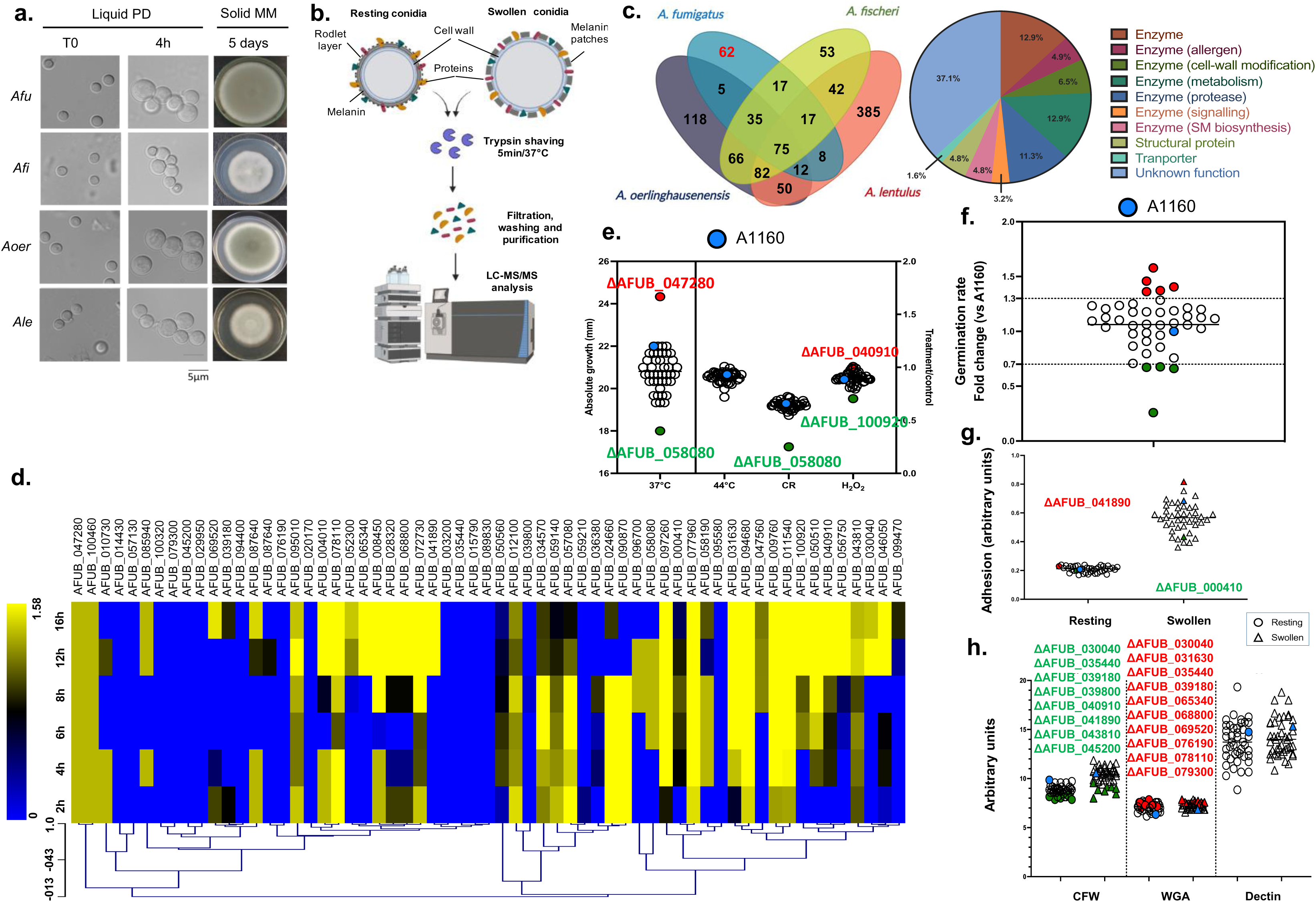
Comparative trypsin-shaving proteomics to study the surface associated proteome “surfome” of *A. fumigatus*. **a.** Representative images of Afu, Afi, Aoer, and Ale conidia at time 0 and 4h post-incubation at 37°C (early germination) in liquid potato dextrose (PD) media and radial growth (10^4^ conidia were displayed at the center of MM plates and incubated for 5 days at 37°C). **b.** Trypsin shaving proteomic analysis workflow. Proteins obtained from trypsin-treated resting and swollen conidia were analyzed by LC–MS/MS. Created with BioRender.com. **c.** Sixty-two unique proteins constitute the *A. fumigatus* conidial surfome. Venn diagram ilustrating the intersection of proteins identified by trypsin-shaving of resting and swollen conidia of *Aspergillus* spp. strains. The conidial surfome of *A. fumigatus* is composed of 62 unique proteins (in red). Functional categorization of the 62 proteins that comprise *A. fumigatus* surfome according to their description in FungiDB. **d.** Heat map of the mRNA accumulation during conidial germination of 62 genes encoding the *A. fumigatus* surfome proteins. The RNAseq database according to Baltussen *et al.* (2023). **e.** Growth phenotypes of the wild-type A1160 strain (in red) and deleted mutants grown for 72 h at 37°C or 44°C in solid MM media and MM supplemented with Congo Red (CR; 10µg/ml) or hydrogen peroxide (H_2_O_2_; 1.5mM). Red circle represents A1163 strain. **f.** Germination rates of the wild-type A1160 strain (in blue) and deleted mutants. Only those hits below or above the 30%-fold change threshold and stastistically different from the parental strain A1160 are displayed in the figure. Blue circle represents A1163 strain. **g.** Adhesion, masured by CV assay, of resting (○) and swollen (△) conidia for all all mutant strains and A1160 (in blue). Only hits that shared the same significant phenotype (lower and higher detection in green and red, respectively) in both stages are highlighted in the figure. Red circle and triangle represent A1163 strain. **h**. Detection of chitin (CFW), N-acetylglucosamine (GlcNAc) (WGA) and β-(1,3)-glucan (Dectin) contents on the conidial surface of all mutant strains and A1160 (in blue) in resting (○) and swollen (△) conidia. Blue circle and triangle represent A1163 strain. Only hits that shared the same significant phenotype (lower and higher detection in green and red, respectively) in both stages are highlighted in the figure.

Comparative analyses of the surface-associated proteome or "surfome" revealed 62 conidial surface proteins were *A. fumigatus*-specific, i.e., only detected in *A. fumigatus* conidia. Fifty-six of the genes encoding these proteins are shared by the majority of 206 *A. fumigatus* isolates, whereas six (AFUB_028320, AFUB_039180, AFUB_045200, AFUB_050510, AFUB_069520, and AFUB_094680) were more variable, belonging to the accessory pangenome (pangenome information from ^18^). In the other three *Aspergillus* species, 53, 118, and 385 conidial proteins were unique to Afi, Aoer, and Ale, respectively. Numerous conidial proteins were also broadly shared; 188 proteins were shared by at least two species, 146 were shared by at least three species, and 75 were shared by all four species (**Figure 2c**).

Functional categorization of the *A. fumigatus*-specific conidial surfome was carried out manually according to the information available in FungiDB database (https://fungidb.org/fungidb/app/). Most genes encode enzymes involved in many biological processes such as cell wall modification, metabolism, cell signaling, and secondary metabolite biosynthesis. Some of these enzymes have previously been identified as allergens and proteases (**Figure 2c**; **Table 1**). Other surface proteins belonged to less enriched functional categories, such as structural proteins and transporters. However, more than a third of the proteins identified as part of the *A. fumigatus*-specific conidial surfome had unknown functions (**Figure 2c**). To further explore the putative function of identified proteins, we predicted the presence of secretion and GPI-anchor peptides using SignalP - 5.0 (https://services.healthtech.dtu.dk/service.php?SignalP-5.0) and PredGPI (http://gpcr2.biocomp.unibo.it/gpipe/index.htm). Thirty-one out of the 62 proteins are predicted to have a signal peptide cleavage site, while only two have high probability (>99%) of harboring a GPI-anchoring signal (**Table 1**). One protein (encoded by Afu3g13755) did not have a homologue in the *A. fumigatus* A1163 background strain and was not examined further. We examined the expression of these 62 genes in a recently published RNAseq dataset that describes gene modulation during conidial germination ^19^ (**Figure 2d**). About 27% of the surfome genes (16 genes) do not change their gene expression during the first 16 h of germination while 33% (21 genes) have increased expression during the 2 to 16 h germination window (**Figure 2d**). About 24% of the surfome genes (15 genes) are expressed late, while 16% (10 genes) are expressed early (**Figure 2d**).

**Table 1.**
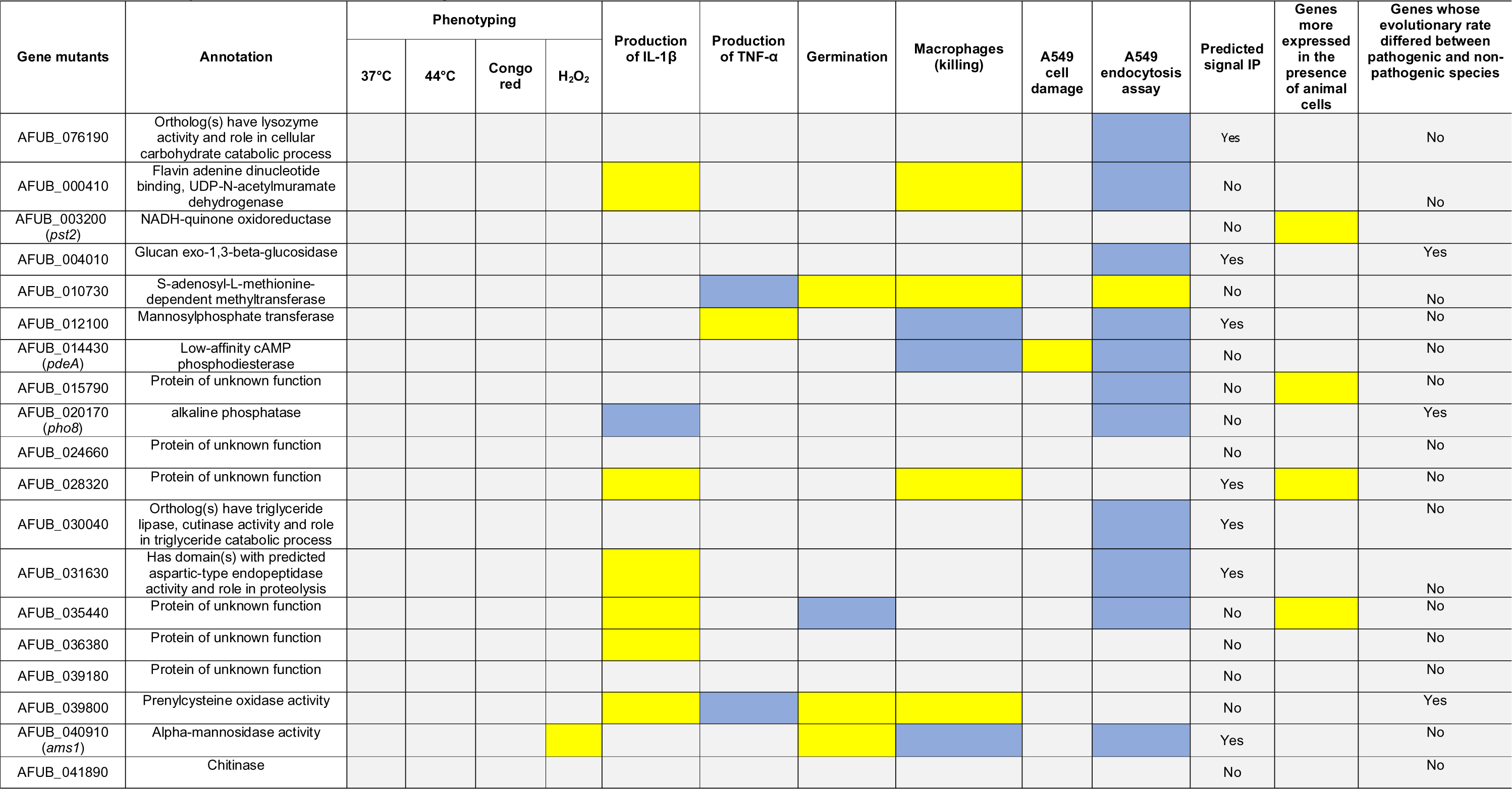
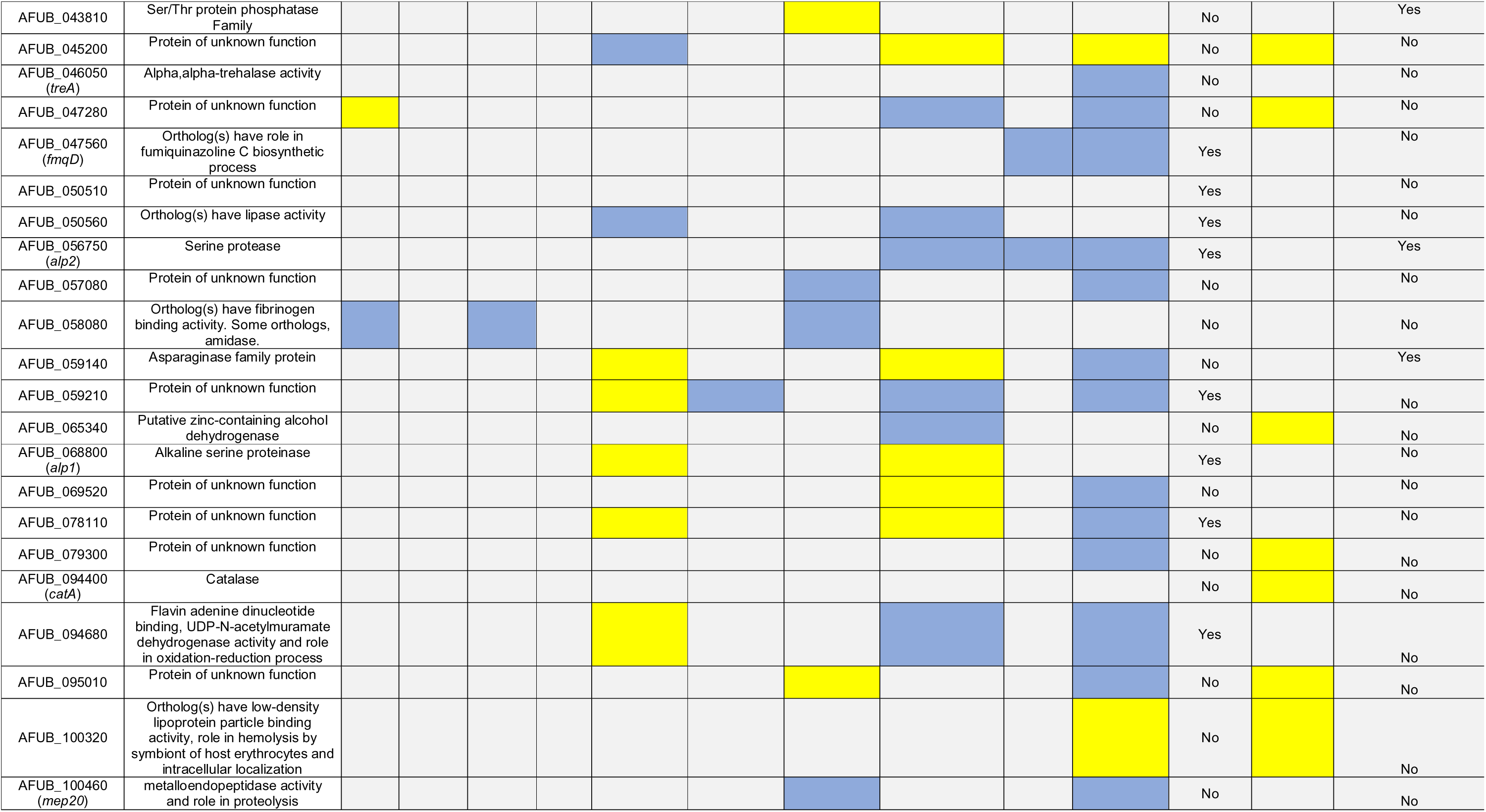
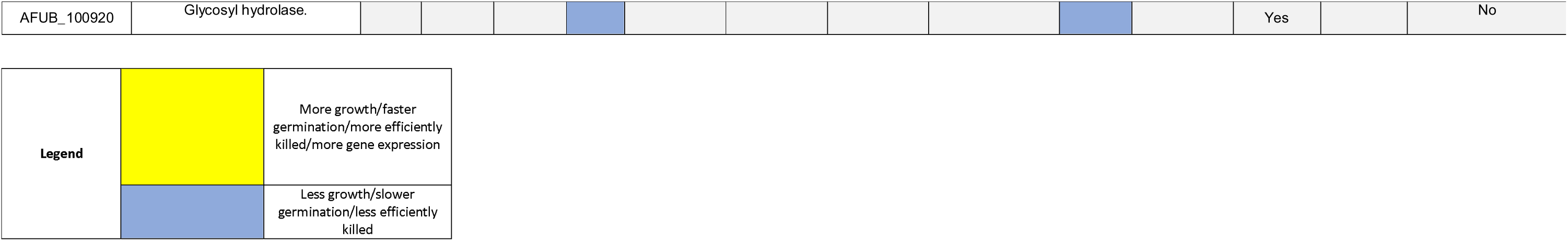
Phenotypes of selected *A. fumigatus* surfome mutants.

Further characterization of *A. fumigatus*-specific conidial surface proteins was carried out using gene-deletion strains and phenotypic assays. To do so, we targeted the construction of homozygous deletion mutants for the 62 genes identified in our previous proteomic analysis following the high-throughput methodology described in ^20^. After three attempts, we constructed deletion mutants for 42 of the 62 *A. fumigatus*-specific genes. It remains to be determined if we were not able to delete the 20 remaining genes because they encode essential proteins. Deletions were PCR validated (primers listed in **Supplementary Table S2**). An exhaustive phenotypic screen was carried out for each deletion mutant to identify the involvement of the *A. fumigatus* surfome in diverse bioprocesses related to virulence (e.g., growth at the temperature of the human body). Growth rates in solid MM at 37°C revealed a defect in growth for ΔAFUB_058080 mutant strain, while ΔAFUB_047280 displayed increased or faster growth compared to the parental A1160 strain. None of the mutants exhibited increased sensitivity to heat (44°C) (**Figure 2e and Table 1**). However, ΔAFUB_058080 was found to be more sensitive than the A1160 wild-type strain to the cell wall stressor Congo Red (CR). Similarly, ΔAFUB_040910 and ΔAFUB_100920 showed decreased and increased susceptibility to oxidative stress, respectively (**Figure 2e and Table 1**). Germination rates for all the mutants and wild-type strain were also determined; most null mutant strains (33/42) behaved similarly to A1160 reference strain. The remaining deletion strains could be differentiated into two groups: i) slow-germinating (below 30% of A1160 germination rate; ΔAFUB_035440, ΔAFUB_057080, ΔAFUB_058080, ΔAFUB_100460) and ii) fast-germinating (above 30% of A1160 germination rate; ΔAFUB_010730, ΔAFUB_039800, ΔAFUB_040910; ΔAFUB_043810, ΔAFUB_095010) (**Figure 2f and Table 1**).

The crystal violet (CV) assay was used as an indirect measurement of the adhesion properties of conidia at both resting and swollen stages for all the strains. Interestingly, significant differences between the wild-type and deletion strains were only observed in two instances: ΔAFUB_000410 displayed reduced adhesion properties in comparison with A1160, while ΔAFUB_041890 showed a higher capacity to adhere to surfaces than the wild-type strain (**Figure 2g and Table 1**).

Abnormalities in conidial cell wall organization were investigated by staining different components of the cell wall. Calcofluor White (CFW) binds and blocks chitin biosynthesis, and FITC-conjugated wheat germ agglutinin (WGA), a lectin that recognizes exposed chitin and chitooligomers were used for staining chitin. Chitooligomer exposure is better quantified using WGA; total cell wall chitooligomer content is better assessed by CFW. CFW revealed a group of 8 strains (ΔAFUB_030040, ΔAFUB_035440, ΔAFUB_039180, ΔAFUB_039800, ΔAFUB_040910, ΔAFUB_041890, ΔAFUB_043810 and ΔAFUB_045200) that exhibited lower chitin exposure than the wild-type strain, and WGA a set of 11 mutants (ΔAFUB_030040, ΔAFUB_031630, ΔAFUB_035440, ΔAFUB_039180, ΔAFUB_065340, ΔAFUB_068800, ΔAFUB_069520, ΔAFUB_076190, ΔAFUB_078110 and ΔAFUB_079300) that showed higher chitin exposure (**Figure 2h and Table 1**). Intriguingly, three of these mutants showed opposing chitin exposure phenotypes (ΔAFUB_030040, ΔAFUB_035440 and ΔAFUB_039180). Using a dectin-based immunofluorescent assay, we did not find any consistent phenotypes for β-D-glucan exposure (**Figure 2h and Table 1**). Finally, only one mutant (ΔAFUB_050510) showed altered hydrophobicity properties when compared with the A1160 wild-type strain (**Supplementary Figure S2**).

Taken together, these results suggest that *A. fumigatus* conidial surface-associated proteins are also involved in the correct assembly of the cell wall and have an important role in combatting stress and adhering to surfaces.

### A subset of *A. fumigatus* surfome proteins is involved in host-pathogen interactions

To understand the role of the *A. fumigatus* surfome in the establishment of infection, we individually assessed the role of each of the 42 *A. fumigatus-specific* surfome proteins in survival and tissue invasion using *in vitro* models of infection. *A. fumigatus* killing was evaluated by the number of CFUs after challenging murine macrophages (Raw 264.7) with conidia from each of the null mutants at 6 h post-infection (hpi), while *A. fumigatus* invasion was determined by differential staining after infecting human lung epithelial cells (A549) with germinating conidia from all deletion strains at 3 hpi. For statistical analyses, we only considered those strains showing a 30% increase or decrease in conidial survival and invasion compared to the parental strain (A1160). Overall, we identified 18 *A. fumigatus* genes encoding surface proteins that contribute to survival to macrophage killing (9 conidial surface null strains were more susceptible to macrophage killing and another 9 were more resistant to being cleared) (**Figure 3a**, **Table 1**). On the other hand, 27 *A. fumigatus* genes encoding for surface proteins were important for epithelial cell invasion (3 mutants were more efficiently taken up by the epithelial cells whilst 24 exhibited defects in cell invasion) (**Figure 3b**, **Table 1**). Although there is no correlation between macrophage survival and epithelial cell invasion (**Figure 3c**), interestingly, 14 *A. fumigatus* surfome proteins were identified to be involved in both killing and invasion (ΔAFUB_000410, ΔAFUB_004010, ΔAFUB_012100, ΔAFUB_014430, ΔAFUB_028320, ΔAFUB_040910, ΔAFUB_045200, ΔAFUB_047280, ΔAFUB_056750, ΔAFUB_059140, ΔAFUB_059210, ΔAFUB_069520, ΔAFUB_078110 and ΔAFUB_094680) (**Figures 3a and 3b**, **Table 1**), half of them with unknown function, suggesting that a subset of *A. fumigatus* conidial surfome proteins identified in our proteomic analysis contributes to triggering immune cell response. The capacity of the null surface protein strains to induce epithelial cell damage was also evaluated with 3 strains producing less damage than the A1160 strain and only one triggering higher cell toxicity (**Figure 3d**, **Table 1**).

**Figure 3.**
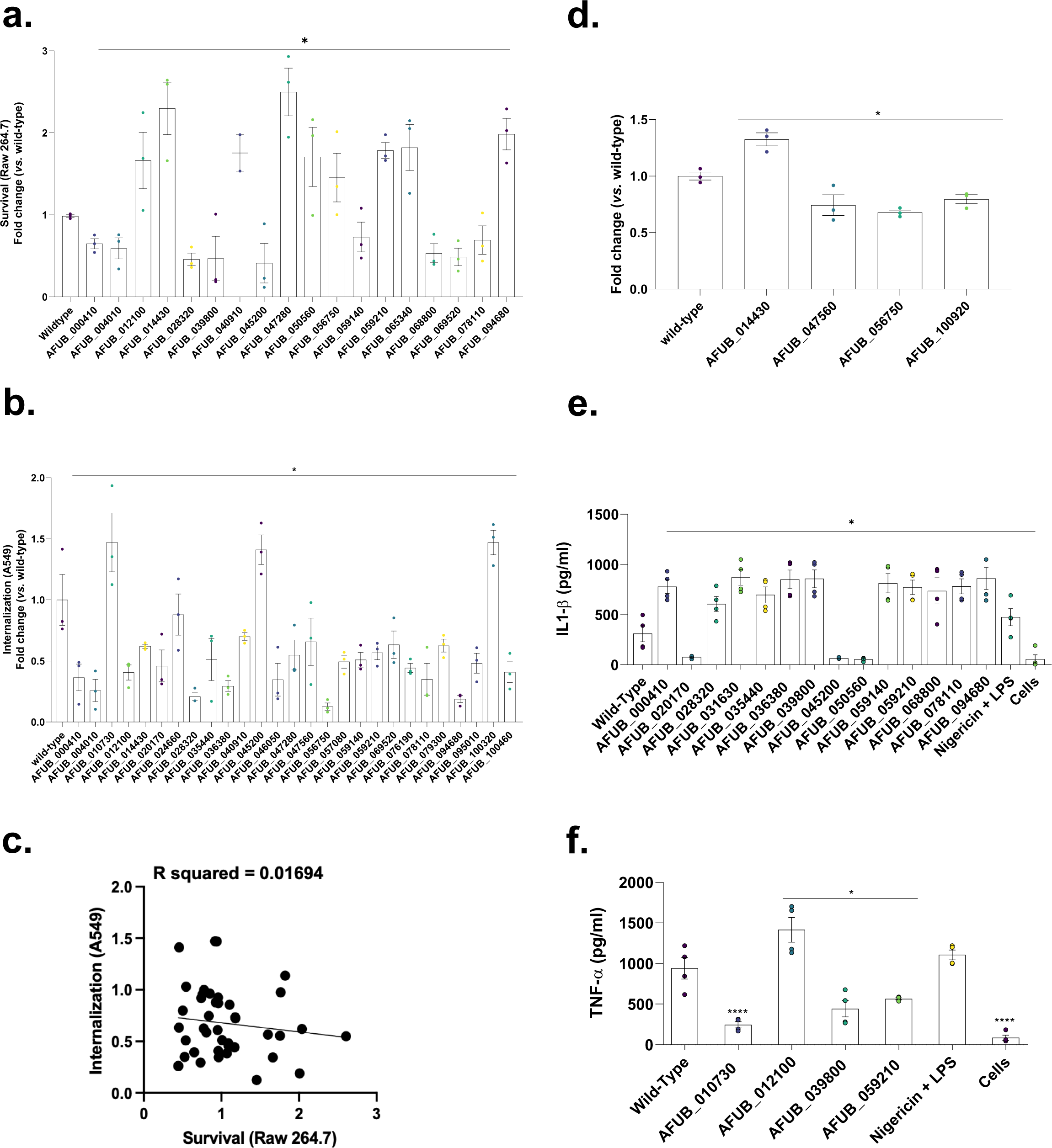
*A. fumigatus* surface-associated proteins are involved in host-pathogen interactions. **a.** Conidia survival of deleted mutants in murine Raw 264.7 macrophages at 6 hpi expressed as fold change against A1160 wild-type strain. **b.** Conidia internalization in A549 epithelial cells at 3 hpi expressed as fold change against A1160 (wild-type) strain. **c**. Simple linear regression models of **a.** and **b.** datasets (no correlation). Thresholds of 30 %-fold change above (red) and below (green) the wild-type strain (A1160) condition were established for statistical significance testing. **d.** Damage to A549 epithelial cells. **e and f.** *A. fumigatus* wild-type and mutant conidia eliciting cytokines IL-1β and TNF-α production by BMDMs. Standard deviations represent averages of results from at least three independent biological repetitions. Statistical analysis was performed using one-tailed, paired t-tests for comparisons to the wild-type strain (A1160) (*p < 0.05).

Our data in **Figure 1e to h** indicate *A. fumigatus* induces a stronger immune response in BMDMs compared to three other closely related species. To investigate whether increased induction of immune response in *A. fumigatus* is driven by specific conidial surface protein(s), we investigated if the absence of these 42 proteins could play a role in cytokine response by determining the IL1-β and TNF-α production in BMDMs (**Figures 3e and 3f and Table 1**). We observed a significantly increased 1L-1β production in 11 mutants (AFUB_000410, AFUB_028320, AFUB_031630, AFUB_035440, AFUB_036380, AFUB_039800, AFUB_059140, AFUB_059210, AFUB_068800, AFUB_078110, and AFUB_094680) and decreased production in 3 mutants (AFUB_020170, AFUB_045200, and AFUB_050560) (**Figure 3f and Table 1**). TNF-α production was significantly increased in AFUB_012100 and decreased in AFUB_010730, AFUB_039800, and AFUB_059210 (**Figure 3f and Table 1**). We also evaluated the BMDMs survival in the presence of these mutants by looking at their lactate dehydrogenase (LDH) activity during infection with the mutants compared to the wild-type strain; only the AFU_065340 (a putative zinc-containing alcohol dehydrogenase) mutant had about 80% inhibited LDH.

Using information about *A. fumigatus* regulatory networks publicly available at the National Center for Biotechnology Information (NCBI) and the clustering online tool FungiExpressZ (https://cparsania.shinyapps.io/FungiExpresZ/), we identified two clusters enriched for RNAseq datasets of *A. fumigatus* exposed to animal cells or invasive aspergillosis conditions (**Figure 4a, highlighted in red circles**). These regulatory datasets were obtained after challenging human dendritic and lung epithelial cells with *A. fumigatus* strains (NCBI Bioprojects PRJEB1583 and PRJNA399754). These two clusters contain 22 genes, 11 of which are also among the 62 *A. fumigatus*-specific genes, suggesting that the protein products of these 11 surfome genes could be important for host-pathogen interactions and the establishment of the infection (**Figures 4b and 4c**, **Table 1**). Other upregulated clusters are mainly related to response to temperature changes, oxidative and osmotic stresses, or nutritional availability, thus illustrating many other functions of *A. fumigatus* conidial surface proteins (**Supplementary Figure S3; Supplementary Table S3**). Furthermore, 7 of the 62 *A. fumigatus*-specific genes were within the 1,700 genes whose evolutionary rate differed between pathogenic and non-pathogenic species from *Aspergillus* section *Fumigati* ^21^, suggesting that they may be associated with the repeated evolution of pathogenicity in *Aspergillus* spp. (**Figure 4d**).

**Figure 4.**
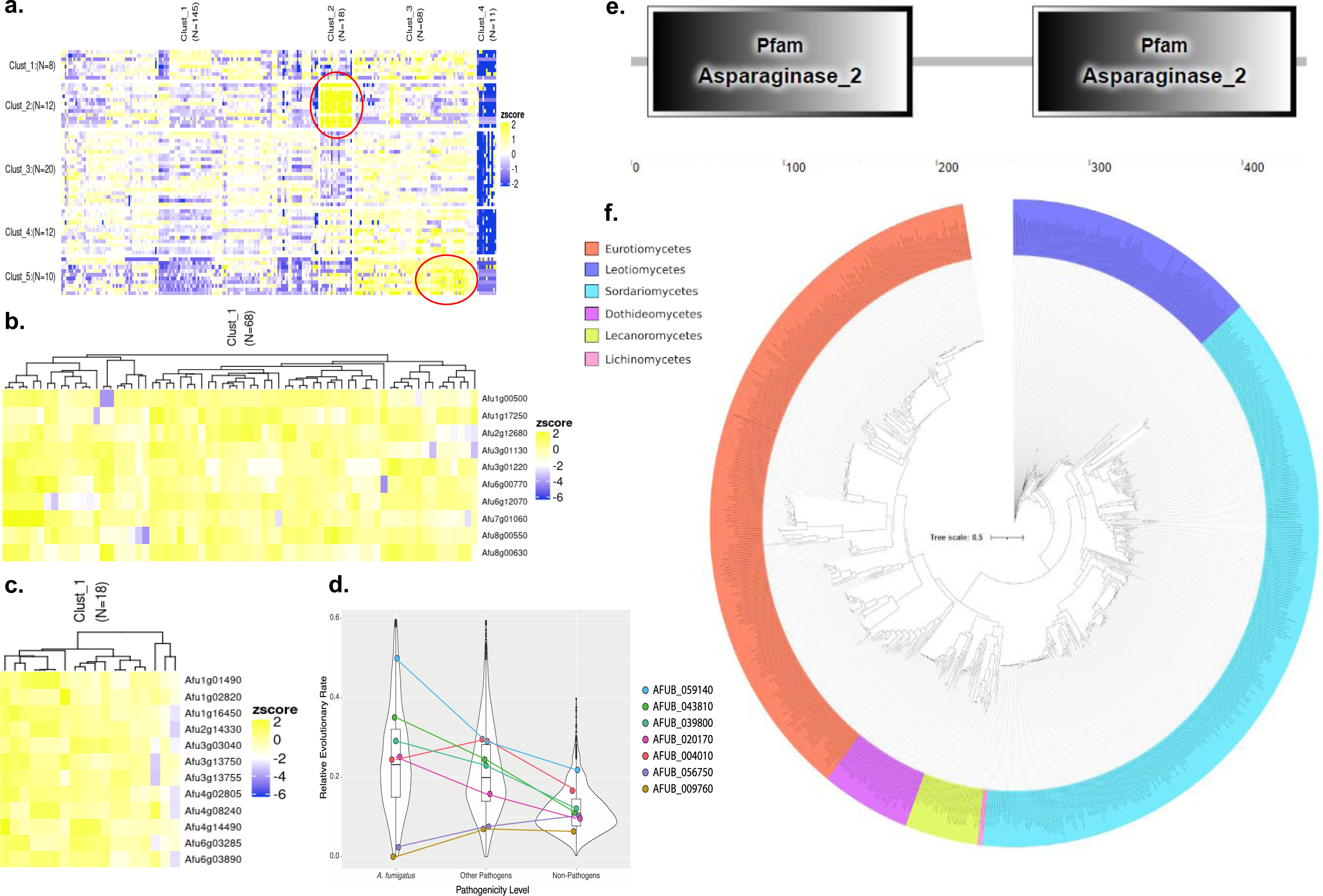
Some of the surfome genes are positively modulated at transcriptional level in the presence of the host cells and a few encoded proteins apparently display faster evolution in pathogenic *Aspergillus spp*. **a.** Heat map clustering of 242 *A. fumigatus* RNAseq datasets available at FungiExpresZ (https://cparsania.shinyapps.io/FungiExpresZ/). **b and c.** Clustering of the two subclusters highlighted in the red circles enriched only for RNAseq datasets of *A. fumigatus* exposed to animal cells or invasive aspergillosis conditions. Heatmap and clustering were generated using the online tool FungiExpressZ (https://cparsania.shinyapps.io/FungiExpresZ/) with the publicly available regulatory information of *A. fumigatus.* d. Violin and box plots showing the ω values (representing the rate of sequence evolution for each gene in every strain) for genes that exhibited different ω values (p-value <0.01) in the three groups of strains (*A. fumigatus*, other pathogens, and non-pathogens). Seven out of the 62 unique genes that encode the *A. fumigatus* surfome exhibit different evolutionary rates in the conserved pathogenicity evolution model. **e**. Protein organization of the AspA according to http://smart.embl-heidelberg.de/. **f.** Phylogenetic distribution of AspA in fungal classes.

These results suggest that some of the proteins specifically identified in the *A. fumigatus* surfome, but which are absent from three other closely related species, are important for mediating host interactions and eliciting cytokines, and are modulated at transcriptional level in the presence of animal cells.

### Characterization of the glycosylasparaginase null mutant

As an initial step to investigate in more detail the surfome mutants, we prioritize the mutants that elicit increased IL-1β production (**Figure 3g and Table 1**) and decided to start by the characterization of ΔAFUB_059140 mutant (here named Δ*aspA*). The *aspA* null mutant has increased killing by macrophages and increased internalization by A549 epithelial cells (**Figures 3a to 3c and Table 1**). We constructed a second independent Δ*aspA* mutant and both independent *aspA* null mutants have the same phenotypes, strongly suggesting that the observed phenotypes are only due to the *aspA* single deletion and not to secondary mutations present in these strains. The *aspA* gene encodes a putative glycosylasparaginase (**Figure 4e**) that catalyzes the hydrolysis of N4-(beta-N-acetyl-D-glucosaminyl)-L-asparagine yielding as products N-acetyl-beta-glucosaminylamine plus L-aspartate cleaving the GlcNAc-Asn bond that links oligosaccharides to asparagine in N-linked glycoproteins, playing a major role in the degradation of glycoproteins (http://pfam-legacy.xfam.org/; PF01112). AspA is distributed among several fungal classes (687 sequences) but enriched in Eurotiomycetes and Sordariomycetes where most of the plant and animal pathogens are present (**Figure 4f**). The Δ*aspA* mutants are more phagocytosed by BMDMs than the wild-type (**Figure 5a**) and have increased percentage of adherent conidia on the BMDMs surface, and decreased cell viability as measured by CFW fluorescence and metabolic activity by MTT, respectively (**Figures 5a to 5c**). The increased expression of 1L-1β in the Δ*aspA* mutants is dependent on their viability since UV-killed conidia have no IL-1β and TNF-α activities comparable to the wild-type strain (**Figure 5d**). Although BMDMs have comparable Reactive Oxygen Species (ROS) production upon 4 and 24 h exposure to either the wild-type or Δ*aspA* mutant, BMDMs have significant increased production of ROS when 8 h in the presence of Δ*aspA* mutant, suggesting that this mutant is either not able to cope with ROS detoxification at this time-point or induces increased ROS production (**Figure 5e**). The lack of *aspA* does not change *A. fumigatus* ability to evade phagolysosome acidification in contrast to the *A. fumigatus* Δ*pksP* mutant (**Figure 5f**). Transwell migration assays show that most of the increased production of IL-1β by *aspA* null mutants need contact with BMDMs (**Figure 5g**). In contrast, TNF-α production is equally dependent on contact for either the wild-type and Δ*aspA* mutants (**Figure 5h**). Taken together, these results suggest that AspA activity is important for BMDMs phagocytosis and viability of *A. fumigatus* conidia, and that AspA can modulate the IL-1β production by direct contact of the conidia with BMDMs.

**Figure 5.**
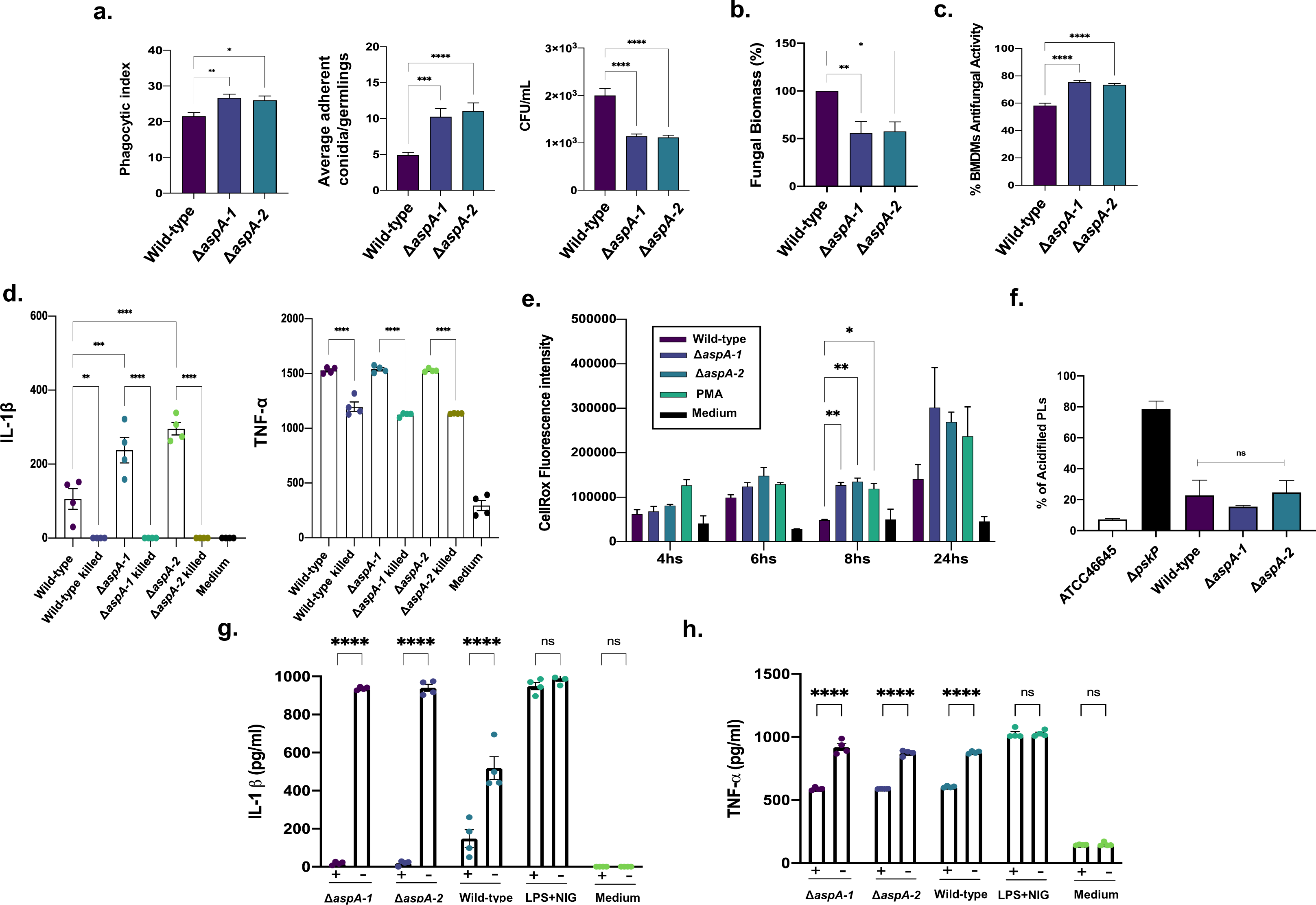
*A. fumigatus* Δ*aspA* mutant has reduced viability in the presence of BMDMs and can modulate IL-1β production. **a.** The Δ*aspA* mutants have increased engulfment and adherence, and reduced viability by the BMDMs when compared to the wild-type strain. **b.** and **c.** Fungal viability upon 24 h exposure to BMDMs measured by MTT and Calcofluor While (CFW) accumulation, respectively. **d.** UV-killed conidia show that Δ*aspA* conidia need to be alive to modulate IL-1β production. **e.** Reactive Oxygen Species (ROS) accumulation upon exposure of the wild-type and Δ*aspA* mutants to BMDMs for 24 h. PMA (phorbol-12-myristate-13-acetate) was used as a positive control for ROS induction. **f.** Percentage of acidifed phagolysosomes (PLs). **g. and h.** Transwell migration assays. Wild-type and Δ*aspA* mutants were exposed to BMDMs and IL-1β and TNF-α production was measured. (+) means the transwell plate and (-) means the control in a ordinary microtiter plate.

### AspA is important for establishment of virulence in an immunocompetent murine model of aspergillosis

*A. fumigatus* Δ*aspA* have the same virulence than the wild-type in a chemotherapeutic murine model but they are less virulent than the wild-type strain in an immunocompetent murine model as measured by fungal burden (**Figures 6a and 6b**). We measured the production of cytokines TNF-α, IL-1β, IL-18, IL-12, INF-□, IL-6, and the chemokine CXCL-1 in the lungs homogenates of the immunocompetent model infected by the wild-type and Δ*aspA* mutants (**Figures 6c to 6i)**. The lack of AspA caused increased production of proinflammatory cytokines and CXCL-1 that acts as a chemoattractant for several immune cells, and it has already been observed as induced upon *A. fumigatus* lung infection ^22^. Our results indicate that AspA is important for attenuating the inflammatory responses and decreasing the neutrophil migration during the early steps of *A. fumigatus* infection in the lungs.

**Figure 6.**
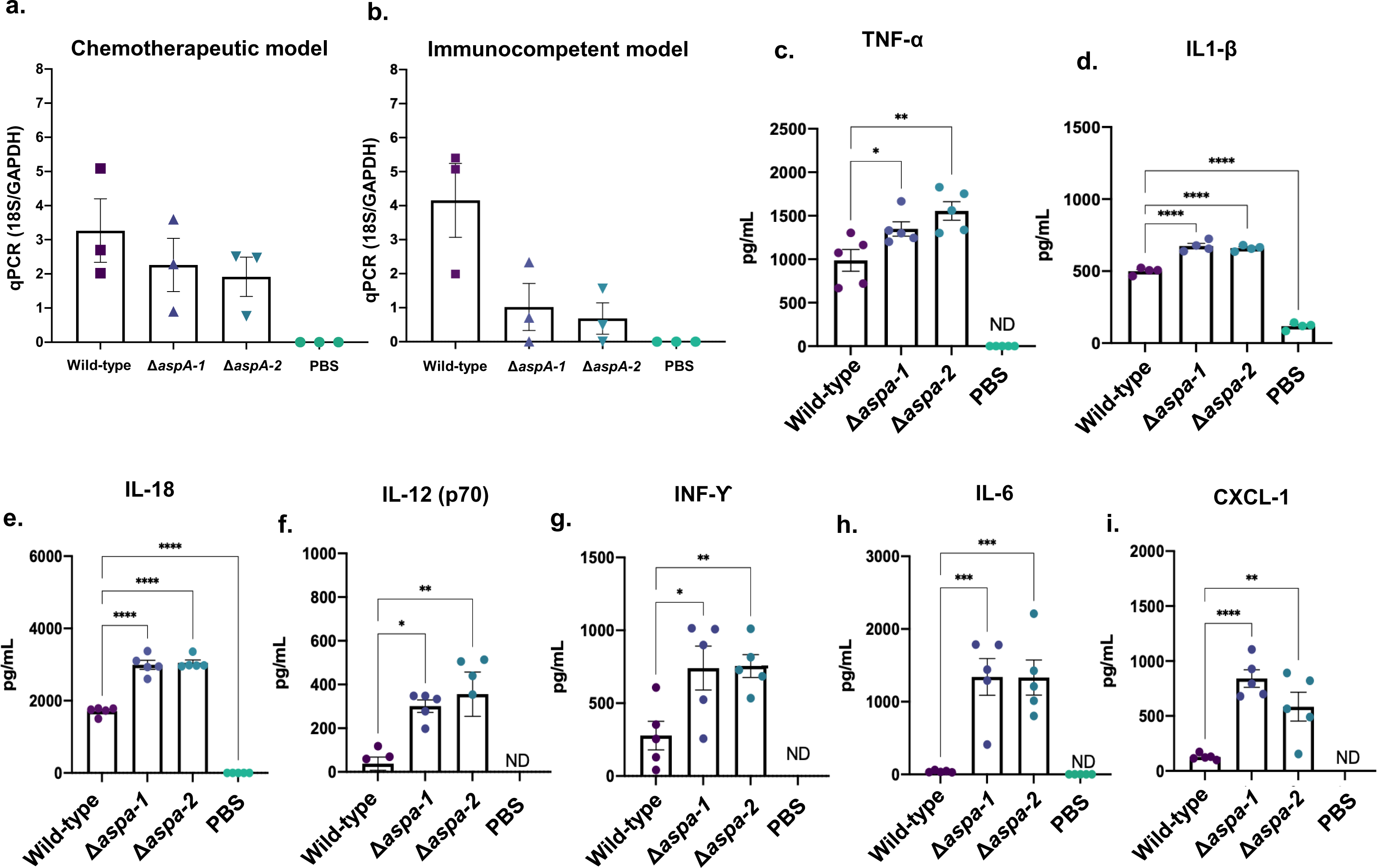
*A. fumigatus* Δ*aspA* mutants have decreased fungal burden in an immunocompetent murine model. **a.** Fungal burden of the wild-type and Δ*aspA* mutants in a murine chemotherapeutic mouse infection model and **b.** immunocompetent model. **c. to h.** Cytokine production (TNF-α, IL-1β, IL-18, IL-12, INF-C, and IL-6) in the immunocompetent mouse model infected by the wild type and Δ*aspA* mutants. **i.** Chemokine production (CXCL-1) in the immunocompetent mouse model infected by the wild type and Δ*aspA* mutants.

### Characterization of the heterologously expressed glycosylasparaginase

We heterologously expressed AspA in *Pichia pastoris* and verified if the enzyme was active by checking its putative asparaginase activity ^23^ (**Supplementary Figure S4**). We were unable to identify asparaginase activity in AspA, and we decided to inactivate the enzyme by boiling it for 5 minutes. Boiling of the AspA decreased IL-1β production by about 50% (**Figure 7a**). Most surprisingly, AspA was also able to induce TNF-α production and AspA inactivation decreased its accumulation by 30% (**Figure 7a**). AspA can modulate IL-1β and TNF-α production by using different AspA concentrations in a linear dose-response relationship (**Figures 7b and 7c**). The cytokine induction by AspA is not related to its glycosylation since AspA exposure to Peptide:N-glycosidase F (PNGase F), which cleaves between the innermost GlcNAc and asparagine residues of high mannose, hybrid, and complex oligosaccharides from N-linked glycoproteins and glycopeptides ^24^, did not abolish this induction (**Supplementary Figure S5**). We constructed an AspA:GFP functional mutant but the low GFP expression did not allow us to perform protein localization studies. As an alternative, we heterologously expressed AspA:GFP in *Pichia pastoris* and used this protein for protein localization studies. Incubation of BMDMs for 24 h with AspA:GFP 0.2 or 2 µg decreased their viability by about 1 and 10%, respectively, while DMSO decreased by 80% (**Figure 7e**); however, we were able to observe fluorescence only with AspA:GFP 2 µg but not 0.2 µg (**Figure 7f**). BMDMs exposed to 2 µg of AspA:GFP for 8 h showed enrichment for vesicles that could be derived from events related to the pinocytosis of the AspA:GFP and/or further fusion of these primary pinocytic vesicles with lysosomes (**Figure 7f**). Weak fluorescence was visualized in about 56% of the cells with possible colocalization with these vesicles when BMDMs were exposed to AspA:GFP for 2 h (**Figure 7f**). There is no autofluorescence when BMDMs were not exposed to AspA:GFP (**Figure 7g**). Next, we investigated if previous (by 8 h) or concomitant exposure of BMDMs to 2µg of AspA and conidia could change the conidial viability (**Figure 7g**). Both treatments increased conidial viability by about 20%, strongly indicating that AspA contributes to conidial viability (**Figure 7g**).

**Figure 7.**
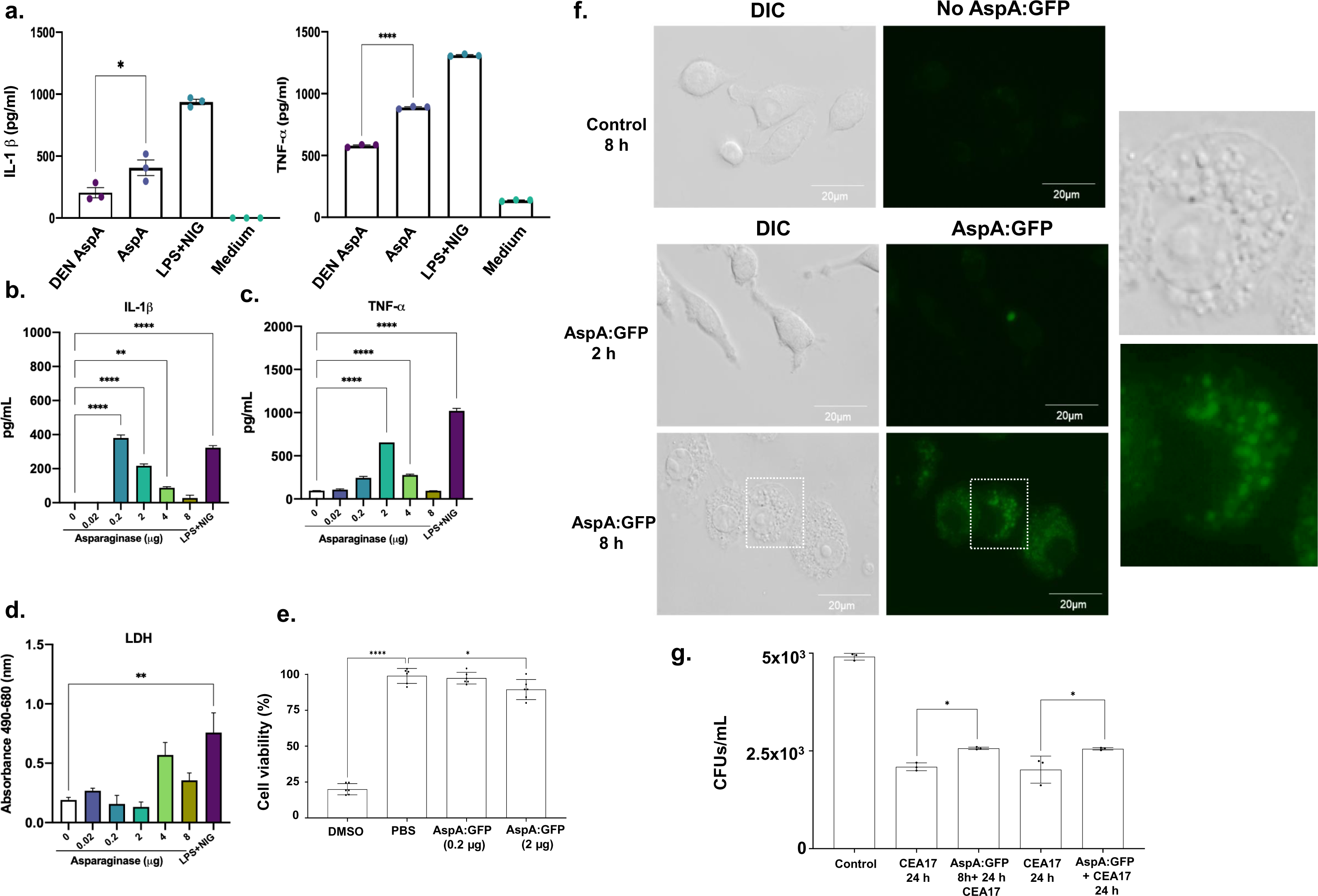
Heterologously expressed AspA can modulate IL-1β and TNF-α levels and suffers pinocytosis by BMDMs. **a.** Denatured AspA has decreased IL-1β and TNF-α in BMDMs. **b. to d.** Increasing concentrations of AspA can modulate IL-1β and TNF-α production by BMDMs and do not affect lactate dehydrogenase (LDH) activity. **e.** Viability of BMDMs exposed to 24 h at 37°C to DMSO, AspA:GFP 0.2 or 2 µg. **f.** BMDMs were not incubated for 8 h, or incubated for 2, and 8 h in the presence of AspA:GFP (2 µg). DIC=Differential Interference Contrast. Inset showing enlarged area hatched in white square. These images are representative of two independent experiments with 40 BMDMs in each experiment. **g.** Conidial viability of BMDMs exposed or not to AspA 2µg for 8h and subsequent addition of conidia or AspA 2µg together with conidia evaluated after 32 and 24 h.

Taken together these results suggest AspA modulates BMDM cytokines and is present in cytoplasmic structures that resemble vesicles and/or lysosomes.

### Proteomic profiling of BMDMs exposed to AspA

To begin investigating which macrophage metabolic pathways are modulated by AspA, we incubated BMDMs in the presence of AspA (0.2 µg) for 24 h, extracted the proteins, and identified them by mass spectrometry (**Supplementary Table S4**). Principal Component Analysis (PCA) and quantitative analysis of these samples showed that BMDM proteins and BMDM proteins exposed to AspA displayed very different distributions (**Figure 8a**). Differentially expressed proteins are defined as those with a minimum of two-fold change in protein abundance (log2FC ≥ 1.0 and ≤ -1.0; FDR of 0.05) when compared to the BMDMs under the equivalent conditions. We observed 101 proteins upregulated and 259 downregulated in the BMDMs exposed to AspA (**Figure 8b and Supplementary Table S4**).

**Figure 8.**
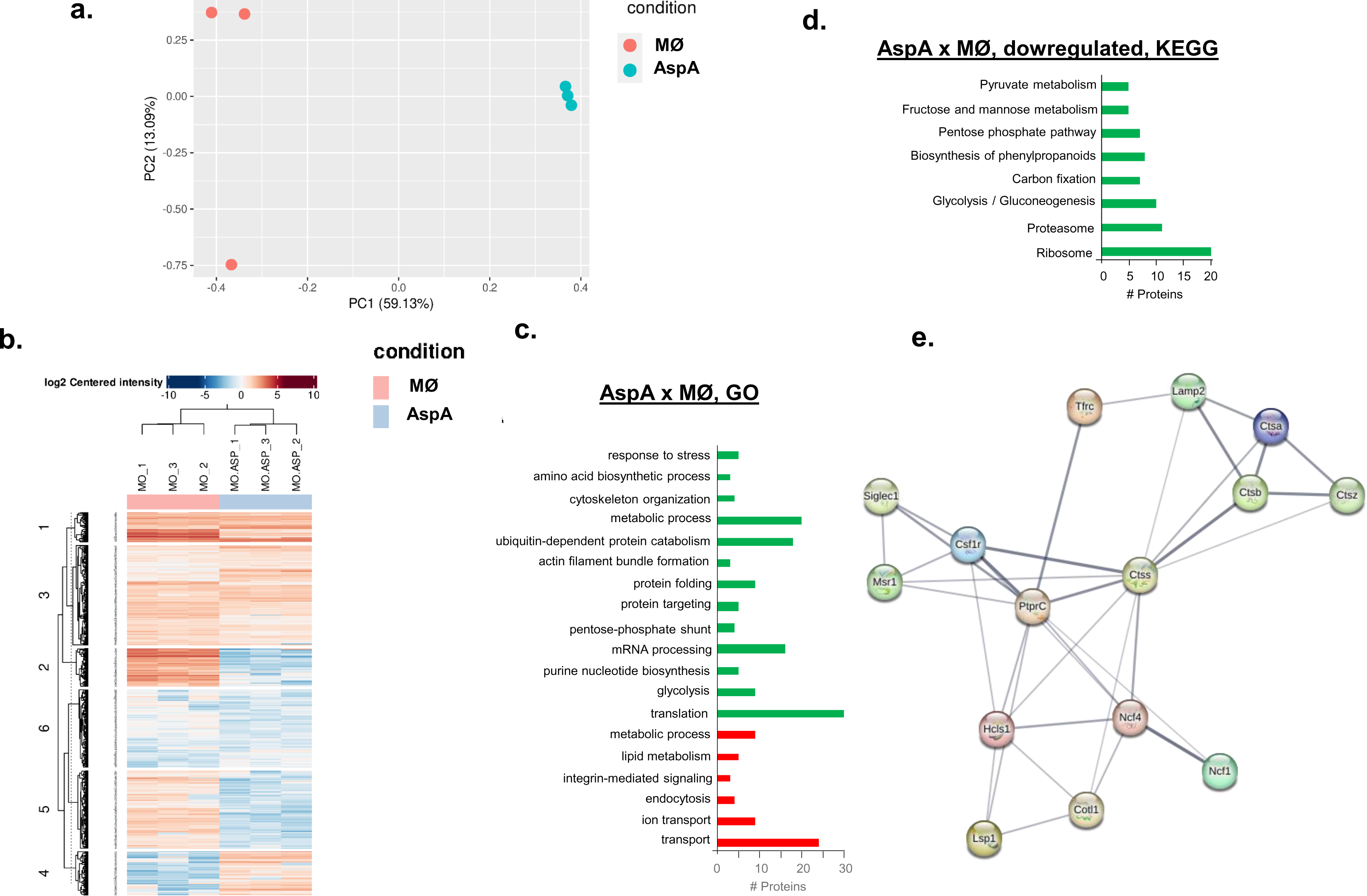
Proteomic profiling of BMDMs exposed to AspA. **a.** PCA distribution of three repetitions of BMDM and BMDM proteins exposed to AspA. **b.** Heat map of protein abundance of BMDM and BMDM exposed to AspA. **c. and d.** Categorization of BMDM and BMDM proteins exposed to AspA. **e.** Mouse functional protein association network based on selected 15 proteins that are modulated in the BMDMs in the presence of AspA. Each node represents a protein that is differentially expressed in the BMDMs upon exposure to AspA. Each edge represents a functional protein association retrieved from the STRING (https://string-db.org) server (medium confidence threshold of 0.4 for the interaction score), and node sizes represent the degree of each node (number of edges connected to the node). Proteins were annotated based on https://www.genecards.org and are: SIGLEC1, Msr1, Csf1r, TfrC, PtprC, Hcls1, Lsp1, Lamp2, Ctss, Ncf4, Colt1, CtsA, CtsB, CtsZ, and Ncf1.

We have used the Reference Database of Immune cells (http://refdic.rcai.riken.jp/document.cgi) for Gene Ontology (GO) and Kyoto Encyclopedia of Genes and Genomes (KEGG) pathway enrichment analysis of distinct biological functions of the shared and specific proteomic differences. The BMDM proteins downregulated in the presence of AspA are involved in (i) cell redox homeostasis, such as glutathione, metabolic process, oxidation reduction, and response to oxidative stress; (ii) carbohydrate metabolism, such as pentose-phosphate shunt, tricarboxylic acid cycle, and glycolysis/gluconeogenesis; (iii) protein synthesis; (iv) pyruvate metabolism and (v) ubiquitin-dependent protein catabolism (**Figures 8c and 8d**). In contrast, protein transport, vesicle-mediated transport, mRNA processing, and metabolic processes are upregulated (**Figure 8c**). We have also visually detected the downregulation of 18 proteins specifically related to the modulation of the innate immune and lysosomal functions (**Table 2 and Supplementary Table S4**), including: (i) Csf1r, macrophage colony-stimulating factor 1 receptor, a tyrosine-protein kinase that acts as a cell surface receptor and plays a role in the regulation of survival, proliferation and differentiation of mononuclear phagocytes, such as macrophages and monocytes (www.uniprot.org/uniprotkb/P09581/entry); (ii) Ncf1, neutrophil cytosol factor 1, required for the activation of the latent NADPH oxidase (www.uniprot.org/uniprotkb/Q09014/entry); (iii) Cotl1, coactosin-like protein that influences leukotrienes synthesis (www.uniprot.org/uniprotkb/Q9CQI6/entry); (iv) Mtdh, protein LYRIC that activates the nuclear factor kappa-B (NF-kappa B) transcription factor (www.uniprot.org/uniprotkb/Q80WJ7/entry), and (v) and a few lysosomal proteases cathepsins; and upregulated, (i) Ptprc, a receptor-type tyrosine-protein phosphatase C that is required for T-cell activation through the antigen receptor (www.uniprot.org/uniprotkb/P06800/entry) and (ii) Ctss, cathepsin S, a key protease responsible for the removal of the invariant chain from MHC class II molecules and MHC class II antigen presentation (www.uniprot.org/uniprotkb/O70370/entry). Examination of the general mouse functional protein association network, retrieved from STRING (https://string-db.org) showed that 15 of these proteins have functional associations, suggesting that AspA impacts specific protein interaction networks (**Figure 8e**).

**Table 2.**
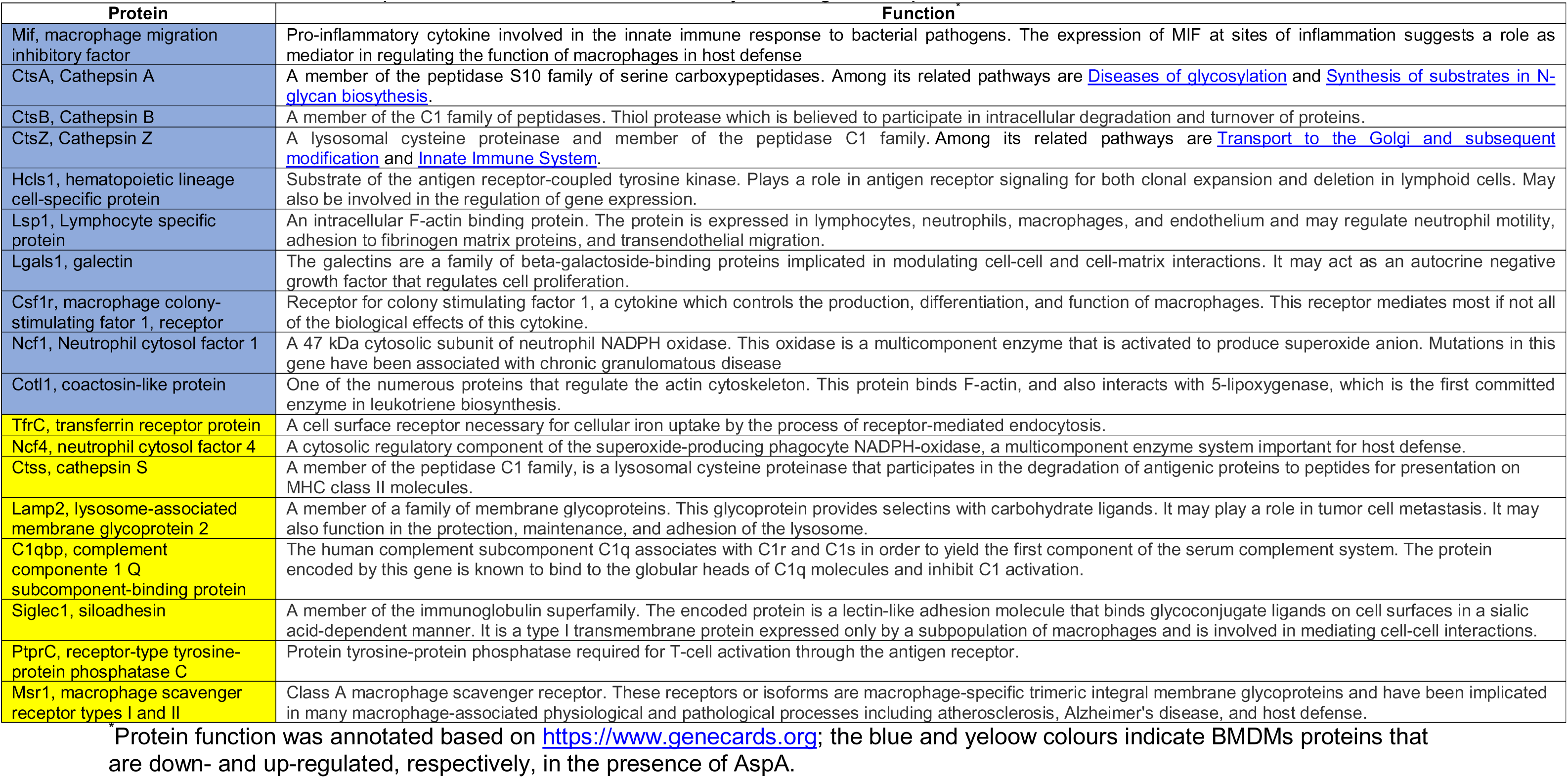
BMDM selected proteins identified as modulated by A. fumigatus AspA.

Taken together, our results suggest that AspA modulates several BMDM metabolic pathways specifically affecting the abundance of proteins important for immune function.

## DISCUSSION

*A. fumigatus* is a prominent human fungal pathogen that infects and frequently kills hundreds of thousands yearly. Most other closely related species, on the other hand, are not pathogenic ^2, 15, 25^. The dozen or so clinically relevant *Aspergillus* species are scattered among hundreds of non-pathogenic species across the *Aspergillus* phylogeny, implying that the capacity to cause illness or pathogenicity has evolved multiple times in this lineage ^2^. The observed pathogenicity spectrum cannot be solely explained by changes in species ecologies or ascertainment bias, suggesting that the repeated evolution of *Aspergillus* pathogenicity has a genetic foundation, at least in part ^2^. *A. fumigatus* conidia are the first fungal structure that encounters the host. In Immunocompetent hosts, conidia are cleared and do not affect host health. However, there are several mechanisms of conidial and germling recognition and evasion based mostly on melanin and polysaccharides, such as β-1,3 glucan, chitin, and galactosaminogalactan present on the conidial surface. Although the importance of protein effectors for pathogenesis and virulence have been extensively described in plant fungal pathogens, there are very few reports about conidial surface proteins or protein effectors in human fungal pathogens. In *A. fumigatus*, these include hydrophobin ^6^, the conidial surface protein *cpcA* ^8^ that prevents immune recognition of β-1,3-glucans, and the HscA protein that anchors human p11 on phagosomal membranes, rewiring the vesicular trafficking to the non-degradative pathway, allowing escape of conidia ^9^. More recently, a *Cryptococcus neoformans* secreted effector that triggers allergic inflammation ^26^ was also identified.

Here, we used a phylogenetic approach to investigate proteins specifically expressed in *A. fumigatus* conidial surface by comparing *A. fumigatus* proteins with two closely non-pathogenic related species, Afi and Aoe, and one more distantly-related pathogenic species, Ale. These surface proteins could be potential surface proteins or protein effectors that modulate different aspects of virulence and pathogenicity or alternatively new genetic determinants important for maintaining the correct conidial structure allowing attachment to host surfaces or preventing immune recognition. Using this approach, we identified 62 *A. fumigatus*-specific conidial surface proteins and were able to delete 42 of them. Some of these proteins affect the correct assembly of the conidial surface components since they have increased or decreased chitin exposure or different hydrophobic properties. However, several of them affect the interaction with the host cells since they modulate survival in the presence of macrophages, invasion and/or damage to epithelial cells, and cytokine production. This collection of genes and mutants provides an opportunity for the characterization of genetic determinants important for the *A. fumigatus*-host interaction.

We focused on one of these genetic determinants, a gene encoding a putative glycosylasparaginase, named AspA. To our knowledge this is the first characterized fungal glycosylasparaginase. Homologues of this gene are not present in *Saccharomyces cerevisiae* and when mutated in humans (AGA gene encodes the glycosylasparaginase) cause the most frequent type of recessively inherited lysosomal storage disease named Aspartylglucosaminuria (AGU) ^27^. AGU is a disorder related to degradation of glycoproteins. Carbohydrate moieties are typically found in glycoproteins and are linked to the protein moiety by an N-glycosidic bond formed by the amino acid L-asparagine and the monosaccharide N-acetylglucosamine. The lysosomal resident enzyme glycosylasparaginase, which cleaves the N-glycosidic link between the L-asparagine and N-acetylglucosamine moieties of GlcNAc-Asn, is lacking in AGU. This enzyme deficiency causes a buildup of undegraded aspartylglucosamine and several other glycoasparagines, i.e., glycoconjugates having a L-asparagine moiety attached to the carbohydrate chain, in the affected person’s tissues and body fluids ^28, 29^. AGU patients have developmental delays, hyperactivity, early growth spurt, inguinal and abdominal hernias, clumsiness, characteristic facial features, recurrent respiratory and ear infections, and chronic arthritis ^30^.

As far as we know there is no previous information about the involvement of glycosylasparaginases in bacterial and fungal pathogenesis. However, there are several reports about possible roles played by asparaginases in bacterial virulence ^31–35^, but not in fungal virulence. Several bacterial virulence factors have evolved to use enzymatic deamidation, i.e., the irreversible conversion of the amino acids glutamine and asparagine to glutamic acid and aspartic acid, respectively ^32^. *Helicobacter pylori* cytotoxic activity was significantly decreased by an asparaginase-deficient strain ^31^ while *Mycobacterium tuberculosis* exploits asparaginase to assimilate nitrogen to produce ammonium and resist acid stress during infection ^33^; *Salmonellae* asparaginases mediate virulence and inhibit T cell responses ^34, 35^. *A. fumigatus* has two putative glycosylasparaginase (AFUB_059140, which is described here, and AFUB_021340, none of them with a putative signal peptide) and two asparaginase encoding-genes (AFUB_003170 and AFUB_051450, with and without a putative signal peptide, respectively). It remains to be investigated if these genes are also important for *A. fumigatus* virulence.

The Δ*aspA* mutants are more phagocytosed and killed than the wild-type, elicit more ROS production and need BMDMs contact to have increased IL-1β production, but have identical percentages of acidification of the phagolysosomes. These results suggest that AspA has a role in earlier or later steps after the formation of phagolysosomes. AspA is important for the establishment of the infection in an immunocompetent mouse model but not in the chemotherapeutic mouse model, and the lack of AspA also plays an important role in modulating proinflammatory cytokines *in vivo*. The lack of AspA causes increased production of proinflammatory cytokines and CXCL-1 that acts as a chemoattractant for several immune cells, especially neutrophils, and it has already been observed as induced upon *A. fumigatus* lung infection ^22^. Our results indicate that AspA is important for modulating the inflammatory response and neutrophils migration upon *A. fumigatus* infection in the lungs.

We also investigated AspA through its heterologous expression and the influence of AspA on cytokine production and macrophage metabolism. Although the human glycosylasparaginase has both deglycosylation and asparaginase activities ^36^, we were not able to detect asparaginase activity in *A. fumigatus* AspA. However, inactivated boiled AspA decreases BMDM cytokine induction, suggesting that it is still active, and this activity is not due to possible glycosylated residues present in AspA since PNGase treatment has not abolished the cytokine induction. AspA has no putative signal peptide, was identified only at the surface of the dormant conidia and does not seem to be secreted. AspA:GFP localizes to BMDMs vesicles that resemble phagolysosomes and/or lysosomes and do not affect BMDM viability. When BMDMs were exposed to AspA, there was a dramatic change in its proteomic profiling, with decreased cell redox and carbohydrate, and pyruvate metabolism, and protein synthesis suggesting that AspA is affecting the BMDMs aerobic metabolism and ROS production, and decreasing the expression of proteins important for ROS production, such as Ncf1, Neutrophil cytosol factor 1, a cytosolic subunit of neutrophil NADPH oxidase, a multicomponent enzyme that is activated to produce superoxide anion. It has been previously shown that patients with hypersensitivity pneumonitis (HP), a rare initial presentation in chronic granulomatous disease (CGD) have mutations in NCF1 and NCF2 genes and are more susceptible to invasive pulmonary *A. fumigatus* infection ^37^. Actually, the Δ*aspA* shows higher induction of ROS in BMDMs. We also observed that BMDMs exposure to AspA decreased the production of several lysosomal proteases, cathepsins, important for protein degradation.

Upon conidial phagocytosis, phagosomes are fused with lysosomes resulting in functional phagolysosomes that are integral part of a degradative pathway that kills and destroys conidia. Is AspA important for escaping phagolysosomal killing? An attractive model for AspA mechanism of action could be that upon conidial phagocytosis, AspA is already interacting with proteins at the macrophage surface deglycosylating asparagine residues and modifying protein activity. Since glycosylasparaginases are described as lysosomal resident proteins, they must be resistant to acid pH (AspA has an isoelectric point of 5.4). Upon conidial phagocytosis and during the formation of phagolysosomes, AspA remains active in the phagolysosome, eventually deglycosylating asparagine residues affecting conidial survival. This apparently has no effect on the percentage of acidification of the phagolysosome since there are no differences between the mutant and wild-type strains. However, it is possible after partial conidial destruction, AspA remains active in the host lysosome affecting the glycosylation metabolism in the cell, and consequently allowing some conidia to survive. *A. fumigatus* conidia can survive for some time in the phagosomes of immune cells and epithelial cells ^38–41^. AspA could affect host’s membrane trafficking system or redirect the conidia-containing phagosomes toward exocytosis and release of conidia. This has already been reported for a novel *A. fumigatus* surface effector, the heat-shock protein HscA that interacts with the human p11 protein, a decisive regulatory node for directing endosomes to different pathways ^9^. Different aspects of our proposed model remain to be investigated.

## METHODS

### Fungal strains

All strains included in this work are listed in **Table 1**. Knockout mutant strains belong to the Manchester Infection Fungal Group (MFIG) collection and were constructed as described in ^20^.

### Surface proteome analysis

Freshly harvested conidia of *A. fumigatus* A1163, *A. oerlinghausenensis* CBS 139183^T^, *A. fischeri* NRRL 181, *and A. lentulus* CNM-CM6069 were cultivated on potato dextrose agar at 37°C for 0 h (resting conidia) or 4 h (swollen conidia). 1 × 10^9^ conidia of each species in three biological replicates were washed twice with 25 mM ammonium bicarbonate (AB) and centrifuged at 1,800 × g for 10 min. Conidia were resuspended in 800 µl of 25 mM AB and incubated with 5 µg trypsin (MS approved, Serva) for 5 min at 37°C. Cell suspensions were immediately passed through 0.2 µm syringe filters (cellulose acetate, Sartorius) to separate conidia from the digestion buffer followed by washing the filters with 200 µl of 25 mM AB. Subsequently, 10 µl of 90 % (v/v) formic acid was added to the cell-free solution to stop the proteolytic digestion. Samples were evaporated to dryness in a vacuum concentrator (Eppendorf), resuspended in 30 µl of 2 % (v/v) acetonitrile (ACN) and 0.05%(v/v) trifluoroacetic acid (TFA), centrifuged for 15 min at 14,000 × g through 10 kDa molecular weight cut-off filters (modified PES, VWR), and transferred into HPLC vials.

LC−MS/MS analysis was performed on an Ultimate 3000 RSLC nano instrument coupled to a QExactive HF mass spectrometer (Thermo Fisher Scientific) as described previously ^12^. Tryptic peptides were trapped for 4 min on an Acclaim Pep Map 100 column (2 cm × 75 μm, 3 μm) at a flow rate of 5 μl/min. Peptides were further separated on an Acclaim Pep Map column (50 cm × 75 μm, 2 μm) using a binary gradient of eluents A (0.1 % (v/v) formic acid in H_2_O) and B (0.1 % (v/v) formic acid in 90:10 (v/v) ACN/H2O): 0-4 min at 4 % B, 5 min at 8 % B, 20 min at 12 % B, 30 min at 18 % B, 40 min at 25 % B, 50 min at 35 % B,57 min at 50 % B, 62−65 min at 96 % B, 65.1−90 min at 4 % B. Positively charged ions were generated by a Nanospray FlexIon Source (Thermo Fisher Scientific) using a stainless steel emitter with 2.2 kV spray voltage. Ions were measured in the Full MS / data-dependent MS2 (Top15) mode. Precursor ions were scanned at *m/z* 300-1500, a mass resolution of 60,000 full width at half maximum (FWHM), an automatic gain control (AGC) target of 1 × 10^6^, and a maximum injection time (maxIT) of 100 ms. Precursor ions were isolated with a width of *m/z* 2.0. Fragment ions generated in the higher-energy collisional dissociation (HCD) cell at 30 % normalized collision energy using N_2_ gas were scanned at 15,000 FWHM, an AGC target of 2×10^5^, and a maxIT of 80 ms. Dynamic exclusion was set to 25 s.

The MS/MS data were searched against the FungiDB databases of *A. fumigatus* Af293 and *A. fischeri* NRRL_181, the NCBI database of *A. lentulus* and the JGI database of *A. oerlinghausenensis* (all downloaded on 2020/10/27) using Proteome Discoverer 2.4 and the algorithms of Mascot 2.4.1, Sequest HT, and MS Amanda 2.0 and MS Fragger 3.0. Two missed cleavages were allowed for the tryptic digestion. The precursor mass tolerance was set to 10 ppm and the fragment mass tolerance was set to 0.02 Da. Modifications were defined as dynamic Met oxidation, protein N-term acetylation and/or loss of methionine. A strict false discovery rate (FDR) < 1% (peptide and protein level) and a search engine score of >30 (Mascot), > 4 (Sequest HT), >300 (MS Amanda) or >8 (MS Fragger) were required for positive protein hits. The Percolator node of PD2.4 and a reverse decoy database was used for qvalue validation of spectral matches. Only rank 1 proteins and peptides of the top scored proteins were counted (**Supplementary Table S1**).

The infected macrophage proteome was performed through lysis in 8M urea in 25 mM AB containing protease inhibitor cocktail using a probe tip sornicator: 40% amplitude for 3 cycles for 10 s and intervals of 10 s. After sonication, samples were centrifuged, the supernatant was transferred to a new tube and quantified using Bradford reagent. Proteins were reduced with 10mM DTT for 30 min at 30 degrees, alkylated with 40mM iodoacetamide for 40 min at room temperature in the dark and overnight digested with trypsin in the ratio 1:50 (enzyme to protein ratio). The tryptic peptides were desalted with Oasis HLB Cartridges (Waters) according to the manufacturer instructions, dried down by speed-vac, and then resuspended in 0.1% formic. LC-MS/MS analysis was performed in an EASY-nLC system (Thermo Scientific) coupled to LTQ-Orbitrap Velos mass spectrometer (Thermo Scientific). Peptides were separated on C18 PicoFrit column (C18 PepMap, 75 µm id□×□10 cm, 3.5 µm particle size, 100 Å pore size; New Objective) using a gradient of A and B buffers (buffer A: 0.1 % formic acid; Buffer B: 95 % CAN, 0.1 % formic acid) at a flow rate of 300nL/min: from 2% to 30% B over 80 min and from 30% to 90% B over 5 min. The LTQ-Orbitrap Velos was operated in positive ion mode with data-dependent acquisition. The full scan was obtained in the Orbitrap with an automatic gain control (AGC) target value of 10e6 ions and a maximum fill time of 500 ms. Each precursor ion scan was acquired at a resolution of 60,000 FWHM in the 400–1500 m/z mass range. Peptide ions were fragmented by CID MS/MS using a normalized collision energy of 35. The 20 most abundant peptides were selected for MS/MS and dynamically excluded during 30 sec. All raw data were accessed in the Xcalibur software (Thermo Scientific). For protein identification, raw data were processed using MaxQuant software version 1.5.3.8. The MS/MS spectra were searched against a protein database composed of Aspergillus fumigatus and Human sequences with the addition of common contaminants with a tolerance level of 4.5 ppm for MS and 0.5 Da for MS/MS. Trypsin was selected as a specific enzyme with a maximum of two missed cleavages. Carbamidomethylation of cysteine (57.021 Da) was set as a fixed modification, and oxidation of methionine (15.994 Da), deamidation NQ (+0.984 Da) and protein N-terminal acetylation (42.010 Da) were set as variable modifications. PSMs, peptides and proteins were accepted at FDR less than 1%.

Multivariate statistical analyses were performed on the protein groups using the label-free quantification (LFQ) results. The analyses were conducted using the LFQ-Analyst web platform ^42^ with default parameters, including a p-value cutoff of 0.05 and a Log2 fold change cutoff of 1.

### BMDMs protein extraction

The BMDMs protein extraction was performed in 8M urea containing protease inhibitor cocktail using a probe tip sornicator: 40% amplitude for 3 cycles for 10 s and intervals of 10 s. After sonication, samples were centrifuged, the supernatant was transferred to a new tube and quantified using Bradford reagent. Proteins were reduced with 10mM DTT, alkylated with 40mM iodoacetamide and overnight digested with trypsin in the ratio 1:50 (enzyme:protein). The tryptic peptides were desalted with Oasis HLB Cartridges, dried down by speed-vac, and then resuspended in 0.1% formic to the LC-MS/MS analysis that was performed in an EASY-nLC system (Thermo Scientific) coupled to LTQ-Orbitrap Velos mass spectrometer (Thermo Scientific). Peptides were separated on C18 PicoFrit column (C18 PepMap, 75 µm idC×C10 cm, 3.5 µm particle size, 100 Å pore size; New Objective) using a gradient of A and B buffers (buffer A: 0.1 % formic acid; Buffer B: 95 % ACN, 0.1 % formic acid) at a flow rate of 300nL/min: from 2% to 30% B over 80 min and from 30% to 90% B over 5 min. The LTQ-Orbitrap Velos was operated in positive ion mode with data-dependent acquisition. The full scan was obtained in the Orbitrap with an automatic gain control (AGC) target value of 10e6 ions and a maximum fill time of 500 ms. Each precursor ion scan was acquired at a resolution of 60,000 FWHM in the 400–1500 m/z mass range. Peptide ions were fragmented by CID MS/MS using a normalized collision energy of 35. The 20 most abundant peptides were selected for MS/MS and dynamically excluded during 30 sec. All raw data were accessed in the Xcalibur software (Thermo Scientific).

For protein identification, raw data were processed using MaxQuant software version 1.5.3.8. The MS/MS spectra were searched against a protein database composed of human sequences with the addition of common contaminants with a tolerance level of 4.5 ppm for MS and 0.5 Da for MS/MS. Trypsin was selected as a specific enzyme with a maximum of two missed cleavages. Carbamidomethylation of cysteine (57.021 Da) was set as a fixed modification, and oxidation of methionine (15.994 Da), deamidation NQ (+0.984 Da) and protein N-terminal acetylation (42.010 Da) were set as variable modifications. Proteins and peptides were accepted at FDR less than 1%.

### Glycosylasparaginase heterologous expression

The synthetic asparaginase gene (AFUB_059140) and its fusion with GFP were cloned in the expression vector pPICZαA (Invitrogen) was synthesized by the company Genescript. The vector has optimized codons for the expression of *Komagataella phaffii* and the α factor secretion signal in the N-terminal portion and Zeocin resistance gene. The plasmid obtained was propagated in competent *Escherichia coli* (DH10β) cells (Thermo Fisher Scientific, Waltham, USA), resistant to zeocin. The purified plasmid was linearized by Anza 24 MssI restriction enzyme (Invitrogen), purified again, and transformed into competent *K. phaffii* (KM71H) by electroporation (1.5 kV, 25 mF and 200 Ω) in a 0.2 µm cuvette using an electroporator (MicroPulser Electroporator, Bio-Rad). The recombinant strain was selected, cultivated and induced to express the rAsparaginase protein following the EasySelect *Pichia Expression* Kit manual (Invitrogen, Waltham, USA). After 120 h of induction with 1% (v/v) methanol, the material was centrifuged (5,000 × *g*, 45 min at 4°C) and the supernatant was filtered through a 0.22 µm filter.

The filtered material was dialyzed against 50 mM Bicine buffer (pH 8.5) and subjected to anion exchange purification. The Mono Q 5/50 GL column (GE Healthcare) was equilibrated with 50 mM Bicine buffer (pH 8.5) and the enzyme eluted with 180 mM NaCl. Protein integrity and sample purity were confirmed by running a 12 % SDS-PAGE ^43^. Purified protein fractions were dialyzed against 1X PBS (pH 7.4) using the Vivaspin 10 kDa centrifuge system (Sartorius). Protein quantification was performed in a spectrophotometer at 280 nm and the assays were performed with the enzyme at a concentration of 5.88 μM.

### AspA phylogenetic analysis

The visualize the distribution of the glycosylasparaginase gene presence among fungi, the AspA amino acid sequence (AFUB_059140) was used for BLAST search ^44^ against the NCBI RefSeq database ^45^. The 687 sequences found were aligned using MAFFT v7.508 ^46^ for the inference of a maximum likelihood phylogenetic tree using IQ-TREE v1.7 ^47^ with the substitution model JTT+I+G4, determined to be the best by the program. The calculated tree was visualized and edited using iTOL v6 ^48^.

### Macrophage Culture

BALB/c bone marrow-derived macrophages (BMDMs) were obtained as previously described ^49^. Briefly, bone marrow cells were cultured for 7–9 days in RPMI 20/30, which consists of RPMI-1640 medium (Gibco, Thermo Fisher Scientific Inc.), supplemented with 20 % (vol/ vol) FBS and 30 % (vol/vol) L-Cell Conditioned Media (LCCM) as a source of macrophage colony-stimulating factor (M-CSF) on non-treated Petri dishes (Optilux - Costar, Corning Inc. Corning, NY). Twenty-four h before experiments, BMDM monolayers were detached using cold phosphate-buffered saline (PBS) (Hyclone, GE Healthcare Inc. South Logan, UT) and cultured, as specified, in RPMI-1640 (Gibco, Thermo Fisher Scientific Inc.) supplemented with 10 % (vol/vol) FBS, 10 U/mL penicillin, and 10 mg/mL streptomycin, (2 mM) L-glutamine, (25 mM) HEPES, pH 7.2 (Gibco, Thermo Fisher Scientific Inc.) at 37°C in 5 % (vol/vol) CO_2_ for the indicated periods.

### Macrophage infection, cytokine and LDH determination

BMDMs were cultured as described before and were seeded at a density of 10^6^ cells/ml in 24-well plates (Greiner Bio-One, Kremsmu□nster, Austria). The cells were challenged with the conidia of different strains at a multiplicity of infection of 1:10 and incubated a 37°C with 5 % (vol/vol) CO_2_ for 24h. BMDMs were also stimulated with different concentrations (mM) of AspA protein (desnaturated or not) by boilling per 10 min 100 degrees Celsius. LPS (standard LPS, *E. coli* 0111: B4; Sigma-Aldrich, 500 ng/mL) plus Nigericin (tlrl-nig, InvivoGen 5 μM/mL) and medium alone were used respectively as the positive and negative controls. Cell culture supernatants were collected and stored at −80°C until they were assayed for TNF-α, IL-1 and LDH release using Mouse DuoSet ELISA kits (R&D Systems, Minneapolis, MN, USA and CyQUANT™ lactate dehydrogenase (LDH) Cytotoxicity Assay (Invitrogen), according to the manufacturer’s instructions. For cytokine determination, plates were analysed by using a microplate reader (Synergy™ HTX Multi-Mode, BioTek) measuring absorbance at 450 nm. Cytokine concentrations were interpolated from a standard curve and statistical significance was determined using an ANOVA (GraphPad Prism 8.0, La Jolla, CA). The level of LDH was determined by measuring absorbance at 490 and 680nm using a microplate reader (Synergy™ HTX Multi-Mode, BioTek). All assays were performed in triplicate in three independent experiments.

### *Aspergillus* growth condition and Fluorescein isothiocyanate (FIT*C)* label

*Aspergillus* strains were cultivated on minimal medium (AMM) agar plates at 37°C for 3 days. Conidia were harvested in sterile water with 0.05 % (vol/vol) Tween20.The resulting suspension was filtered through two layers of gauze (Miracloth, Calbiochem). FITC-labelling of conidia was performed with 0.1 mg/ml FITC (Sigma) in 0.1 M Na_2_CO_3_ at 37 °C for 30 min. Labelled conidia were washed three times with PBS, 0.1% (vol/vol) Tween20. The conidia concentration was determined using a hemocytometer.

### Phagocytosis and adhesion assays

BMDMs were cultivated in Dulbecco’s Modified Eagle Medium (DMEM) supplemented with 10% (vol/vol) heat-inactivated fetal calf serum, 2mM glutamine and penicillin-streptomycin. For infection experiments, macrophages were seeded on glass cover slips in 24 well plates at a density of 5×10^5^ cells per well and allowed to grow adherently overnight. Following washing with prewarmed medium, FITC-labelled conidia were added at a multiplicity of infection of 10. The infection experiment was synchronized for 30 min at 4°C. Unbound conidia were removed by washing with pre-warmed medium and phagocytosis was initiated by shifting the coincubation to 37°C in a humidified CO_2_ incubator. After 1 h the phagocytosis was stopped by washing with ice-cold PBS. Labelling of extracellular conidia was performed by incubation with PBS, 0.25 mg/ml calcofluor white (Sigma) for 30 min at 4°C to avoid further. The cells were washed twice with PBS and fixed with 3.7 % (vol/vol) formaldehyde/PBS for 15 min followed by two washes with PBS. Microscopic photographs were taken on a Zeiss microscope. For statistical reproducibility two biological replicates and in each case two technical replicates were made and analyzed for each strain. The phagocytic index was enumerated by counting 100 macrophages per cover slip from duplicate wells. Phagocytic index was calculated by the average number of conidia that had been phagocytosed for each macrophage.

### Macrophage killing assay

BMDMs were seeded at a density of 10^6^ cells/ml in 24-well plates (Corning Costar) and were challenged with conidia at a multiplicity of infection of 1:10 and incubated a 37°C with 5 % (vol/vol) CO_2_ for 24h. After incubation media was removed the cells were washed with ice-cold PBS and finally 2ml of sterile water was added to the wells. A P1000 tip was then used to scrape away the cell monolayer and the cell suspension was collected. This suspension was then diluted 1:1000 and 100 μl was plated on Sabouraud agar before the plates were incubated a 37°C overnight and the colonies were counted. 50 μl of the inoculum adjusted to 10^3^/ml was also plated on SAB agar to correct CFU counts. The CFU/ml for each sample was calculated and compared to the A1160 wild-type strain.

### Acidification of phagolysosomes

Acidification of phagolysosomes was essentially determined as previously described ^17^. In brief, RAW 264.7 murine macrophages were incubated in Dulbecco’s Modified Eagle Medium (DMEM, Gibco) with 27.5 mg/mL gentamicin sulfate (Gibco), 10 % (vol/vol) fetal bovine serum (FBS, GE Healthcare Life Sciences), and 1% (wt/vol) ultraglutamine (Gibco) at 37°C and 5 % (vol/vol) CO_2_ in humidified incubator. Cells were pre-stained with 50 nM LysoTracker Red DND-99 (Invitrogen) for 1 h before infection. Conidia were stained with 0.1 mg/mL Calcofluor White (CFW, Sigma Aldrich) for 10-15 min before infection. 1 x 10^5^ cells per well were added to 8 well ibidi slides (ibidi GmbH) and infected with an MOI of 3 with CFW stained conidia for 2 h at 37°C and 5 % (vol/vol) CO_2_ in a humidified incubator. Uptake of conidia was synchronized by centrifugation at 100 x g for 5 min. After infection, cells were washed with PBS and fixed using 3.7% formaldehyde in PBS for 10 min, followed by a washing step with PBS. Slides were directly imaged using a Zeiss LSM 780 microscope and analyzed using Zeiss ZEN software. At least, 100 conidia were counted for each biological replicate. Clear red signals surrounding conidia within phagosomes were evaluated as acidified conidia. Three independent experiments were performed.

### BMDMs antifungal activity

BMDMs were isolated as described before and 5×10^5^ cells were seeded in 96 round bottom wells in complete DMEM (DMEM (Gibco), 10 % (vol/vol) FBS (Gibco), 1 % (wt/vol) penicillin/streptomycin (Sigma), 1 % HEPES (Gibco). After adhesion the BMDMs were infected with 5×10^6^ conidia (MOI 1:10). Upon 24 h of co-incubation the cells were lysed with sterile water and conidia were stained for 5 min with 50 μL of a filter sterilized 1mg/mL Calcofluor white solution (Sigma-Aldrich). Conidia were washed with sterile water and the staining was quantified using a fluorometer at 360 nm excitation and 440 nm emission.

XTT assay was performed as described ^50, 51^. Briefly, 5 × 10^5^ BMDMs were plated in 96-well and then infected with 5 × 10^6^ spores in a final volume of 100 μl DMEM media. Control wells contained spores but no BMDMs or only BMDMs. Following 2 hr incubation, the macrophages were subjected to hypotonic lysis by three gentle washes with distilled water followed by a 30 min incubation with distilled water at 37°C. Supernatants then were removed, with great care taken not to remove the spores. DMEM media without Phenol Red containing 400 μg/ml of XTT and 50 μg/ml of Coenzyme Q, were added, and the wells were incubated for 2 hr at 37°C. The OD_450_ and OD_650_ were then measured, and data were expressed as the percent of antifungal activity according to the published formula ^50^.

### Cytokine and chemokine quantification

For lung homogenates, the lungs of all experimental groups were homogenized in PBS supplemented with Complete Mini protease inhibitor tablets (Roche), clarified by centrifugation, and stored at −80°C. A panel of cytokines and chemokines were quantified by ELISA (R&D Systems) according to the manufacturer’s instructions.

### Detection of reactive oxygen species (ROS)

In 96-well tissue culture (TC)-treated dark clear bottom plates (Sigma-Aldrich), 5×10^5^ BMDMs per 100 μL were infected with 5×10^6^ conidia followed by 4, 8, and 24 h of incubation at 37 ^0^C. The cells were treated with the ROS detection dye CellROX^®^ Reagent (ThermoFisher) at a final concentration of 5 μM for 30 min at 37^°^C. ROS detection was performed using a fluorescent plate reader (Thermo Fisher), with an excitation of 485 nm and an emission of 520 nm. BMDMs stimulated with 50 nM phorbol-12-myristate-13-acetate (PMA) (Sigma-Aldrich) for 20 min at 37^°^C were used as positive control.

### Transwell Assay

BMDMs were seeded at a density of 10^6^ cells/ml in 24-well plates (Corning Costar) and the assay was performed as the addition each well containing a transwell insert (0.2-μm filter; Corning) containing medium alone, Δ*aspA-1*, Δ*aspA-2* or wild-type conidia at a multiplicity of infection of 1:10 and incubated a 37°C with 5 % (vol/vol) CO_2_ for 24h. After incubation cell culture supernatants were collected and stored at −80°C until they were assayed for TNF-α, IL-1 production. In the transwell system the small and soluble compounds are able to migrate between the upper transwell and lower chamber, whereas spores are prevented from moving between the chambers.

### Epithelial cell invasion assay

The capacity of the various strains to invade the A549 cells was determined using previously described methods with some modification ^52, 53^. The *A. fumigatus* wild-type and mutant strains were grown on Sabouraud dextrose agar (Difco) at 37°C for 7 d prior to use. Conidia were harvested with phosphate-buffered saline (PBS) containing 0.1 % (vol/vol) Tween 80 (Sigma-Aldrich) and enumerated with a hemacytometer. The A549 pulmonary epithelial cell line (American Type Culture Collection) was cultured in F12 medium (American Type Culture Collection) containing 10% (vol/vol) fetal bovine serum (Gemini Bio-Products), and 2 mM L-glutamine with penicillin and streptomycin (Irvine Scientific) in 5% CO_2_ at 37°C. Before the assay, 2 x 10^5^ A549 cells were cultured in 24-well tissue culture plates containing fibronectin coated circular glass coverslips in each well for overnight. *A. fumigatus* conidia were pre-germinated in Sabouraud dextrose broth (Difco) at 37°C for 5.5 h, counted and suspended in F12k medium. Next, 10^5^ germlings of each strain in 1 ml F12 K medium were added to A549 cells that had been grown to confluency on the glass coverslips. After incubation for 3 h, the cells were rinsed with 1 ml HBSS in a standardized manner and then fixed with 4 % (vol/vol) paraformaldehyde. The noninternalized portions of the organisms were stained with a polyclonal rabbit anti-*A. fumigatus* primary antibody (Meridian Life Science, Inc.) followed by an AlexaFluor 568-labeled secondary antibody (Life Technologies). After the coverslips were mounted inverted on microscope slides, they were viewed by epifluorescence. The number of cell-associated organisms was determined by counting the number of red fluoresced organisms per high-powered field (HPF). The number of endocytosed organisms was determined by subtracting the number of non-internalized organisms (staining of entire germlings) from the number of cell-associated organisms. At least 100 organisms per coverslip were scored and each strain was tested in triplicate.

### Epithelial cell damage assay

To evaluate the capacity of various strains to damage A549 epithelial cells, we used our standard ^51^Cr release assay ^53–55^. *A. fumigatus* conidia and A549 were prepared as described before. The A549 cells were grown to confluency in a 24-well tissue culture plate and then loaded with ^51^Cr overnight. The following day, the cells were rinsed twice with HBSS to remove the unincorporated ^51^Cr and 5 x 10^5^ conidia in 1 ml of F12K medium of each strain were added to triplicate wells. After 16 h of incubation, 500 μl of medium above the cells were collected and transferred to glass test tube A. Next, the remaining medium in the wells was collected and placed in glass test tube B. After lysing the A549 cells with 6 N NaOH, the lysate was collected and the wells were rinsed twice with RadiacWash (Biodex Medical Systems). The lysate and rinses were added to test tube B. The amount of ^51^Cr in the medium and the cell lysate was measured using a gamma counter. The spontaneous release of ^51^Cr was determined using uninfected A549 cells that were processed in parallel. The specific release of ^51^Cr was calculated using our previous described formula. Each experiment was performed in triplicate. Mutants that caused less damage to the A549 cells than the control strain were tested at least one more time to verify the results.

### Animal survival curves and fungal burden

Inbred female mice (BALB/c or C57BL/6 strains; body weight, 20–22 g) were housed in vented cages containing five animals. Mice were immunosuppressed with cyclophosphamide (150 mg/kg of body weight), which was administered intraperitoneally on days - 4, -1 and 2 prior to and post infection (infection day is “day 0”). Hydrocortisonacetate (200 mg/kg body weight) was injected subcutaneously on day -3. Mice (10 mice per group) were anesthetized by halothane inhalation and infected by intranasal instillation of 20 µL of 1.0 x 10^5^ conidia of *A. fumigatus* wild-type or mutant strains, *A. lentulus*, *A. oerlinghausenensis*, or *A. fischeri* (the viability of the administered inoculum was determined by incubating a serial dilution of the conidia on MM medium, at 37°C). As a negative control, a group of 10 mice received PBS only. Animals were sacrificed 15 days post-infection. To investigate fungal burden in murine lungs, mice are immunosuppressed as described previously or not, and mice were intranasally inoculated with 1 x 10^6^ conidia/20 µl of suspension for the chemotherapeutic murine model and with 5 x 10^8^ conidia/20 µl for the immunocompetent murine model. Animals were sacrificed 48 h post-infection, and the lungs were harvested and immediately frozen in liquid nitrogen. DNA was extracted via the phenol/chloroform method and 400 mg of total DNA from each sample were used for quantitative PCRs using primers to amplify the 18S rRNA region of *A. fumigatus* and an intronic region of mouse GAPDH (glyceraldehyde-3-phosphate dehydrogenase). Six-point standard curves were calculated using serial dilutions of gDNA from *A. fumigatus* strain and the uninfected mouse lung. Fungal and mouse DNA quantities were obtained from the threshold cycle (CT) values from the appropriate standard curves. The qPCR analysis was performed using the SYBR green PCR master mix kit (Applied Biosystems) in the ABI 7500 Fast real-time PCR system (Applied Biosystems, Foster City, CA, USA).

The principles that guide our studies are based on the Declaration of Animal Rights ratified by UNESCO on January 27, 1978 in its 8th and 14th articles. All protocols adopted in this study were approved by the local ethics committee for animal experiments from the University of São Paulo, Campus of Ribeirão Preto (Permit Number: 08.1.1277.53.6; Studies on the interaction of *A. fumigatus* with animals). Groups of five animals were housed in individually ventilated cages and were cared for in strict accordance with the principles outlined by the Brazilian College of Animal Experimentation (COBEA) and Guiding Principles for Research Involving Animals and Human Beings, American Physiological Society. All efforts were made to minimize suffering. Animals were clinically monitored at least twice daily and humanely sacrificed if moribund (defined by lethargy, dyspnea, hypothermia and weight loss). All stressed animals were sacrificed by cervical dislocation.

### Phenotypic assay

Plates containing solid MM were inoculated with 10^4^ spores per strain and left to grow for 72 h at 37 or 44°C. When required MM was supplemented with Congo Red (10 µg/ml) or H_2_O_2_ (1.5 mM). All radial growths were expressed as ratios, dividing colony radial diameter of growth in the stress condition by colony radial diameter in the control (MM at 37°C) condition.

### Germination assay

The germination process was followed for the A1160 strain over a 16 h time period under a Nikon TI microscope equipped with a 37°C incubator and a 40× objective, with 1 picture captured every 30 min by using NIS-Elements 4.0 (Nikon) software. Cell Counter plug-in of ImageJ platform (http://rsb.info.nih.gov/ij/index.html) was employed to differentially count resting or swollen conidia versus germinated conidia. A time point was selected (7h) in which A1160 achieved 50-60% of germination and the knockout mutant strains were assayed for the same time point included A1160 as a control. 10^4^ conidia were inoculated into 200 μl of liquid MM in 96-well plates and each strain was tested in technical duplicates.

### Staining for cell surface components

Cell wall surface polysaccharide staining was performed as described previously ^56^. Briefly, 10^4^ spores for each deletion mutant and A1160 strains were inoculated in 200µl of MM liquid medium and incubated for 4 h at 37°C (swollen conidia) and 4°C (resting conidia) before the culture medium was removed and conidia were UV irradiated (600,000mJ). For dectin staining, 200 µl of a blocking solution (2% [wt/vol] goat serum, 1 % [wt/vol] bovine serum albumin [BSA], 0.1 % [vol/vol] Triton X-100, 0.05 % [vol/vol] Tween 20, 0.05 % [vol/vol] sodium azide, and 0.01 M PBS) was added to each working well. Samples were incubated for 30 min at room temperature (RT) and 0.2 µg/ml of Fc-h-dectin-hFc (Invivogen) was added to the UV-irradiated conidia and incubated for 1 h at RT, followed by the addition of 1:1,000 DyLight 594-conjugated goat anti-human IgG1 (Abcam) for 1 h at RT. Conidia were then washed with phosphate-buffered saline (PBS), and fluorescence was read at 587-nm excitation and 615-nm emission. For chitin staining, 200µl of a PBS solution with 10mg/ml of calcofluor white (CFW) was added to the UV-irradiated conidia, which were incubated for 5 min at RT and washed with PBS before fluorescence was read at 380-nm excitation and 450-nm emission. For N-acetyl-D-glucosamine (GlcNAc) staining, 200 µl of PBS supplemented with 0.1 mg/ml of wheat germ agglutinin (WGA) lectin (lectin-FITC L4895; Sigma) was added to the UV-irradiated germlings for 1 h at RT. Germlings were washed with PBS, and fluorescence was read at 492 nm excitation and 517 nm emission. All experiments were performed using at least 4 repetitions, and fluorescence was read in a microtiter plate reader (Synergy HTX Multimode Reader; Agilent Biotek or EnSpire Multimode Plate Reader; Perkin Elmer).

### Fungal adhesion assay

In order to determine the adhesion capacity of conidia from deleted mutant and A1160 strains, 10^4^ conidia were inoculated into 200 ul of MM liquid medium in a 96-well polystyrene microtiter plate. Following an initial adherence phase of 4 h during static incubation at 37°C (swollen conidia) or 4°C (resting conidia), unbound conidia were washed with sterile PBS. Fresh MM liquid medium was added to the adhered conidia, and static submerged cultures were grown for up to 24 h at 37°C. Subsequently, the plate was washed exhaustively with PBS prior to incubation with 200 ul 0.5 % (wt/vol) crystal violet solution for 5 min at room temperature. The stained mycelia were then exhaustively washed with sterile water and air dried. Finally, the crystal violet was eluted from the wells using 100 % ethanol, and the absorbance was measured at 590 nm in a Synergy HTX Multimode Reader (Agilent Biotek).

### Assessment of conidial hydrophobic properties

The distribution of the conidia on a water-oil interface was performed by the addition of 1x10^8^ conidia of the wild-type and each deleted strain in a solution containing water and tributyrin (1:1 [vol/vol]). The mixture was vortexed during 1 min and set to allow the conidia to disperse for 1 h. The distribution of the conidia for each mutant were then visually compared to the A1160 wild-type strain.

### Statistical analysis

Grouped column plots with standard deviation error bars were used for representations of data. For comparisons with data from wild-type or control conditions, we performed one-tailed, paired *t* tests or one-way analysis of variance (ANOVA). All statistical analyses and graphics building were performed by using GraphPad Prism 5.00 (GraphPad Software).

### Data Availability

The mass spectrometry proteomics data have been deposited to the ProteomeXchange Consortium via the PRIDE partner repository ^57^ with the dataset identifiers PXD031199 (Username; Password: HjEpfE78) and PXD044190 (Username; Password: WNvL2PfA).

## Acknowledgements

We thank the Fundação de Amparo à Pesquisa do Estado de São Paulo (FAPESP) grants numbers 2021/04977-5 (G.H.G.), 2018/18257-1 (GP), 2018/15549-1 (GP), 2020/04923-0 (GP) and the Conselho Nacional de Desenvolvimento Científico e Tecnológico (CNPq) grant numbers 301058/2019-9 and 404735/2018-5 (G.H.G.), both from Brazil, and the National Institutes of Health/National Institute of Allergy and Infectious Diseases grant R01 AI153356 from the USA (to A.R. and G.H.G.). This work was supported by the Deutsche Forschungsgemeinschaft (DFG, German Research Foundation) Collaborative Research Centre/Transregio FungiNet 124 ‘Pathogentic fungi and their human host: Networks of Interaction’ (project number 210879364; projects A1 and Z2).

