## Supplementary Figure S1 for "A phylogenetic approach to explore the *Aspergillus fumigatus* conidial surface-associated proteome and its role in pathogenesis"

### Supplementary Figure 1

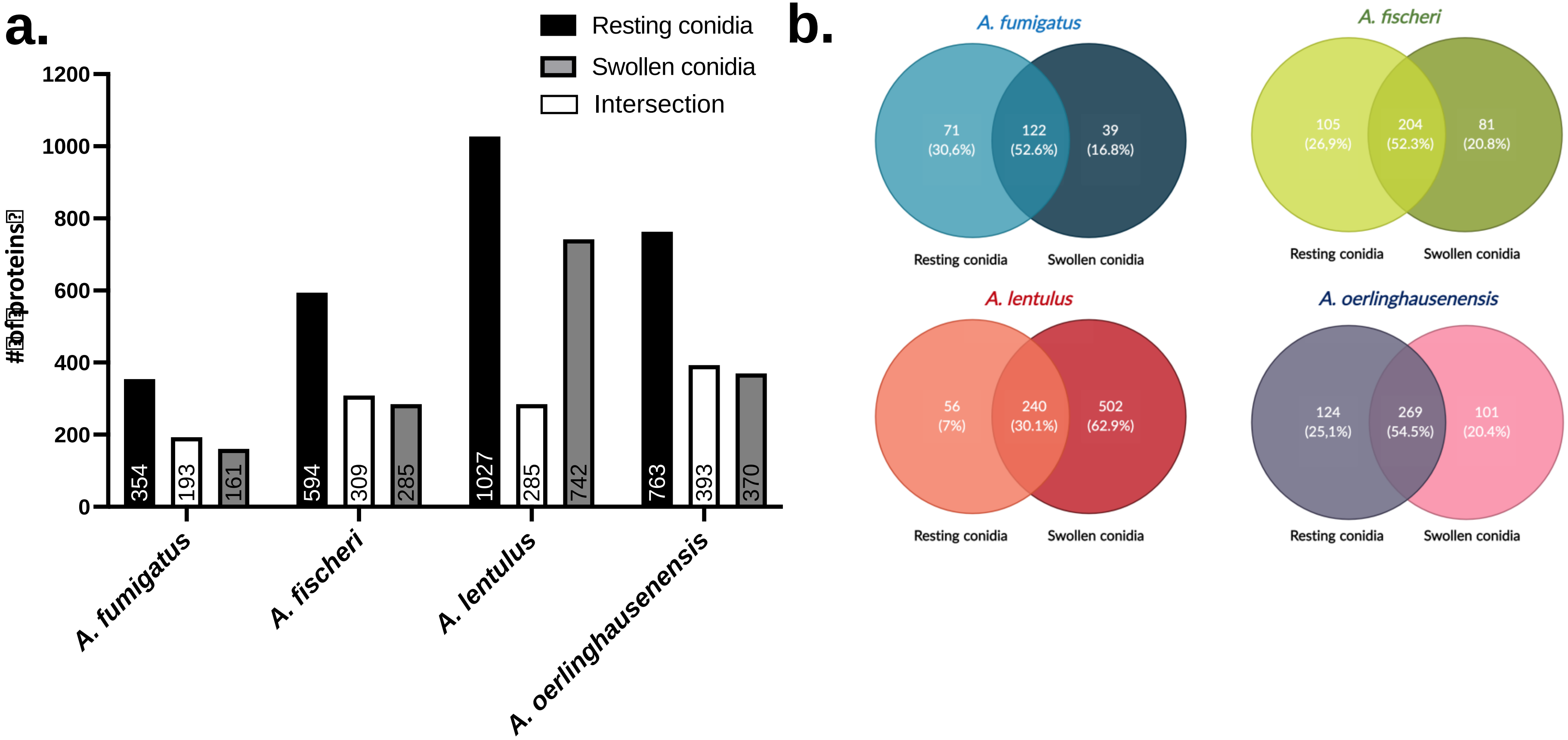


**Supplementary Figure S1. a.** Number (#) of proteins (Found in 3 Biol. Repl.; (PSMs/AAs)*(Cov/100) > 0.001)) identify by trypsin-shaving proteomics in resting and swolllen conidia of the four *Aspergillus* species included in this work. **b.** Venn diagrams ilustrating the intersection of proteins identified by trypsin shaving of resting and swollen conidia of *Aspergillus* spp. strains.
