## Supplementary Figure S2 for "A phylogenetic approach to explore the *Aspergillus fumigatus* conidial surface-associated proteome and its role in pathogenesis"

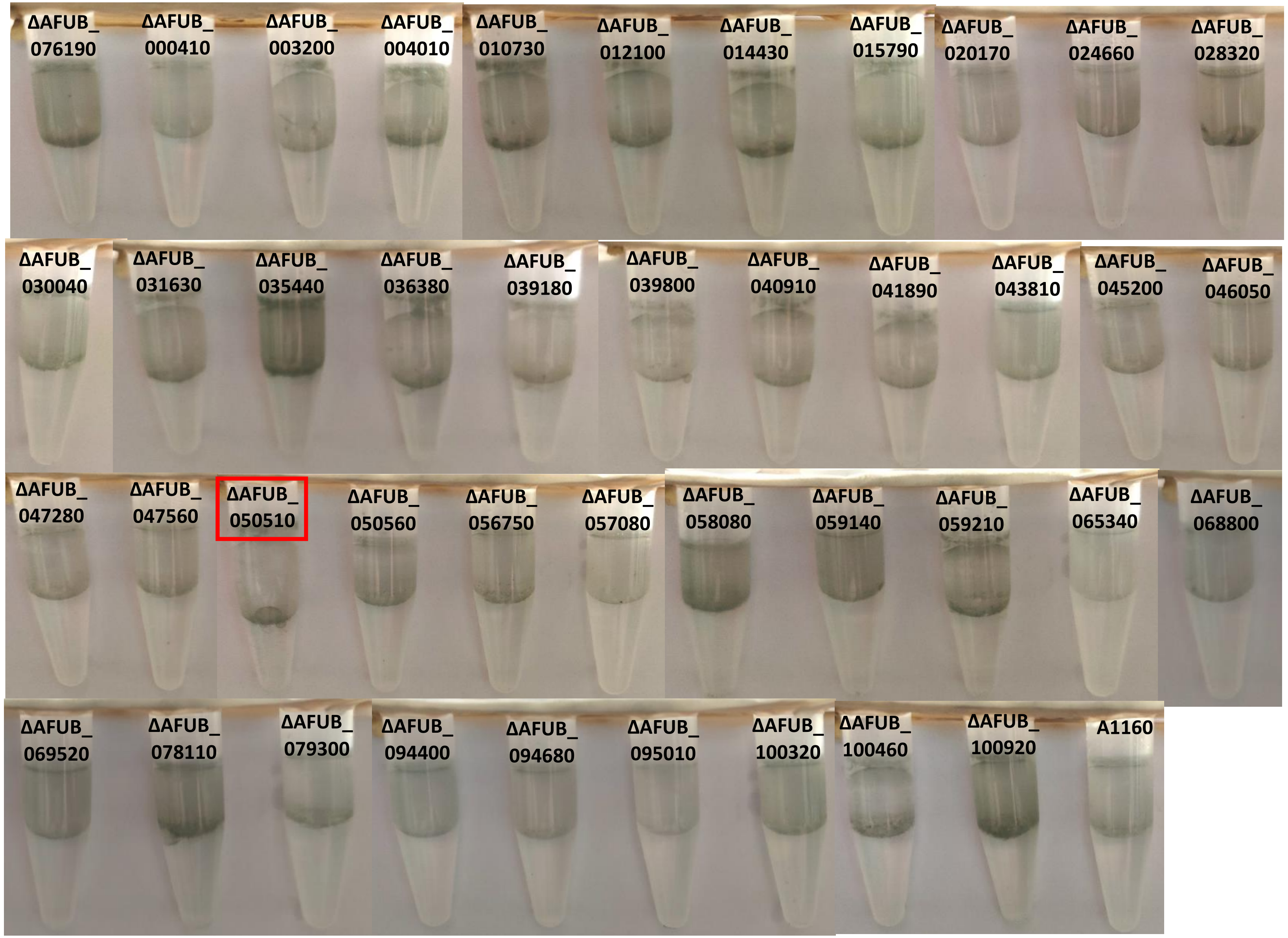


**Supplementary Figure S2.** Distribution of condida of A1160 and deleted mutant strains in a 1:1 water-oil (tributyrin) interface.
