## Supplementary Figure S3 for "A phylogenetic approach to explore the *Aspergillus fumigatus* conidial surface-associated proteome and its role in pathogenesis"

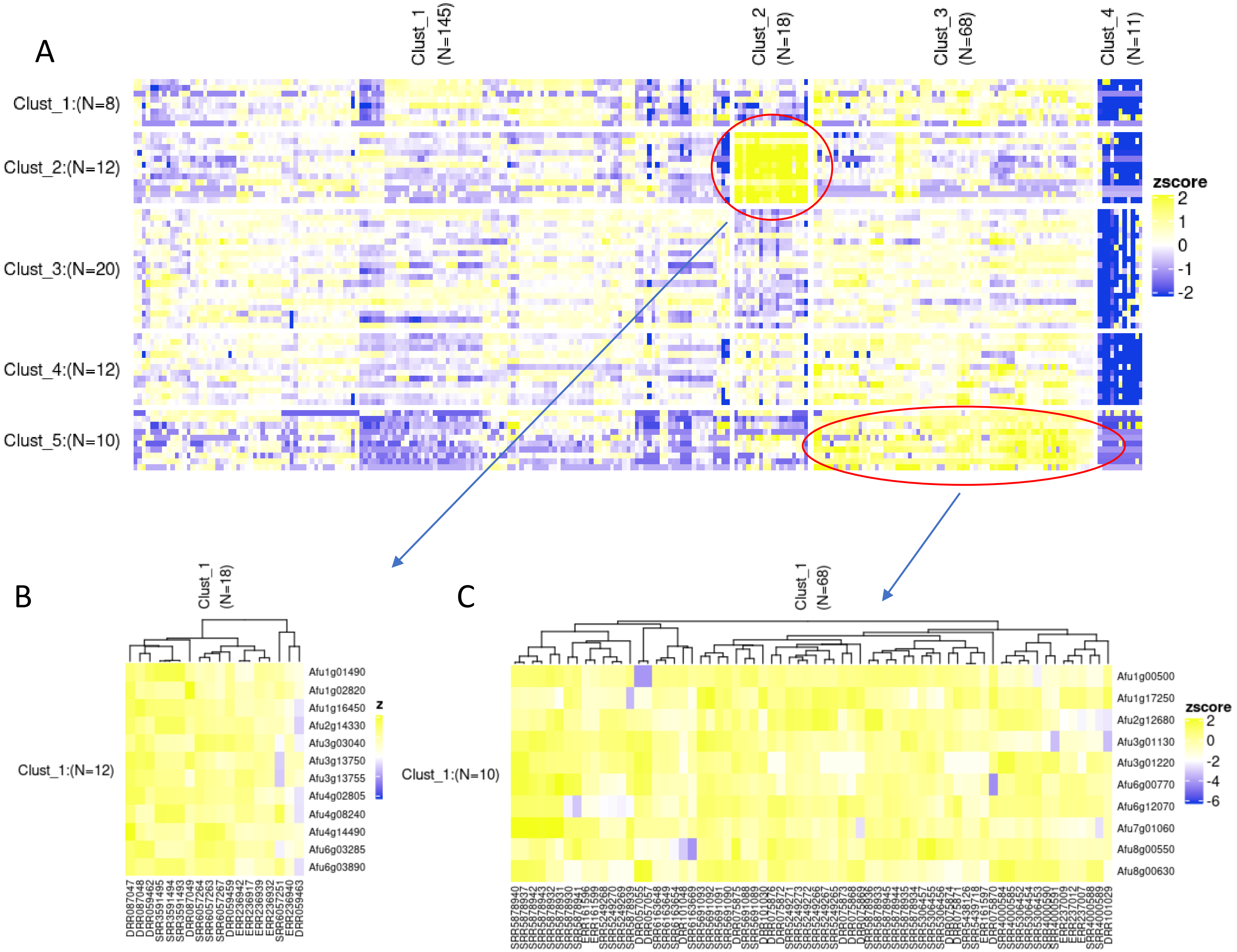


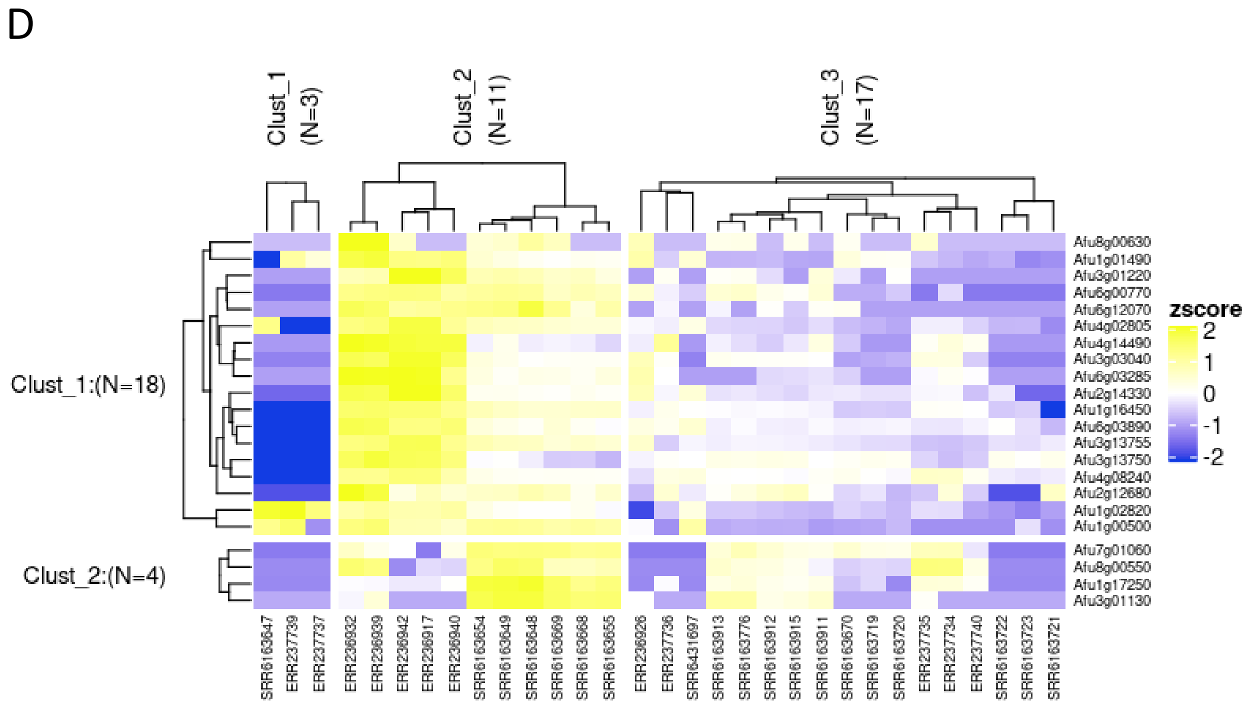


Figure: Clustering of expression data.
A. Expression data of all tested genes on FungiExpresZ entire dataset;
B. Zomm in Cluster 2 super-regulated genes;
C. Zoom in cluster 5 super-regulated genes;
D. Expression of genes from cluter 2 and 5 on animal model experiments.

### Experiments on cluster D.

| column_labels | bio_project | study_title |
|---|---|---|
| ERR236932 | PRJEB1583 | RNA-Seq of healthy human dendritic cells after challenge with four different fungal pathogens, and of four fungal pathogens after challenge with healthy human dendritic cells, SYBARIS project. |
| ERR236939 | PRJEB1583 | RNA-Seq of healthy human dendritic cells after challenge with four different fungal pathogens, and of four fungal pathogens after challenge with healthy human dendritic cells, SYBARIS project. |
| ERR236942 | PRJEB1583 | RNA-Seq of healthy human dendritic cells after challenge with four different fungal pathogens, and of four fungal pathogens after challenge with healthy human dendritic cells, SYBARIS project. |
| ERR236917 | PRJEB1583 | RNA-Seq of healthy human dendritic cells after challenge with four different fungal pathogens, and of four fungal pathogens after challenge with healthy human dendritic cells, SYBARIS project. |
| ERR236940 | PRJEB1583 | RNA-Seq of healthy human dendritic cells after challenge with four different fungal pathogens, and of four fungal pathogens after challenge with healthy human dendritic cells, SYBARIS project. |
| SRR6163654 | PRJNA399754 | In vitro infection of A549 cells with Aspergillus fumigatus |
| SRR6163649 | PRJNA399754 | In vitro infection of A549 cells with Aspergillus fumigatus |
| SRR6163648 | PRJNA399754 | In vitro infection of A549 cells with Aspergillus fumigatus |
| SRR6163669 | PRJNA399754 | In vitro infection of A549 cells with Aspergillus fumigatus |
| SRR6163668 | PRJNA399754 | In vitro infection of A549 cells with Aspergillus fumigatus |
| SRR6163655 | PRJNA399754 | In vitro infection of A549 cells with Aspergillus fumigatus |
| ERR236926 | PRJEB1583 | RNA-Seq of healthy human dendritic cells after challenge with four different fungal pathogens, and of four fungal pathogens after challenge with healthy human dendritic cells, SYBARIS project. |
| ERR237736 | PRJEB2987 | Aspergillus_fumigatus_and_invasive_aspergillosis |
| SRR6431697 | PRJNA399754 | In vitro infection of A549 cells with Aspergillus fumigatus |
| SRR6163913 | PRJNA399754 | In vitro infection of A549 cells with Aspergillus fumigatus |
| SRR6163776 | PRJNA399754 | In vitro infection of A549 cells with Aspergillus fumigatus |
| SRR6163912 | PRJNA399754 | In vitro infection of A549 cells with Aspergillus fumigatus |
| SRR6163915 | PRJNA399754 | In vitro infection of A549 cells with Aspergillus fumigatus |
| SRR6163911 | PRJNA399754 | In vitro infection of A549 cells with Aspergillus fumigatus |
| SRR6163670 | PRJNA399754 | In vitro infection of A549 cells with Aspergillus fumigatus |
| SRR6163719 | PRJNA399754 | In vitro infection of A549 cells with Aspergillus fumigatus |
| SRR6163720 | PRJNA399754 | In vitro infection of A549 cells with Aspergillus fumigatus |
| ERR237735 | PRJEB2987 | Aspergillus_fumigatus_and_invasive_aspergillosis |
| ERR237734 | PRJEB2987 | Aspergillus_fumigatus_and_invasive_aspergillosis |
| ERR237740 | PRJEB2987 | Aspergillus_fumigatus_and_invasive_aspergillosis |
| SRR6163722 | PRJNA399754 | In vitro infection of A549 cells with Aspergillus fumigatus |
| SRR6163723 | PRJNA399754 | In vitro infection of A549 cells with Aspergillus fumigatus |
| SRR6163721 | PRJNA399754 | In vitro infection of A549 cells with Aspergillus fumigatus |
| SRR6163647 | PRJNA399754 | In vitro infection of A549 cells with Aspergillus fumigatus |
| ERR237739 | PRJEB2987 | Aspergillus_fumigatus_and_invasive_aspergillosis |
| ERR237737 | PRJEB2987 | Aspergillus_fumigatus_and_invasive_aspergillosis |

### Experiments on cluster C.

| column_labels | bio_project | study_title |
|---|---|---|
| SRR5878940 | PRJNA396210 | Aspergillus fumigatus delbrlA, delabaA, delwetA, and wild-type RNA-seq |
| SRR5878937 | PRJNA396210 | Aspergillus fumigatus delbrlA, delabaA, delwetA, and wild-type RNA-seq |
| SRR5878942 | PRJNA396210 | Aspergillus fumigatus delbrlA, delabaA, delwetA, and wild-type RNA-seq |
| SRR5878943 | PRJNA396210 | Aspergillus fumigatus delbrlA, delabaA, delwetA, and wild-type RNA-seq |
| SRR5878932 | PRJNA396210 | Aspergillus fumigatus delbrlA, delabaA, delwetA, and wild-type RNA-seq |
| SRR5878931 | PRJNA396210 | Aspergillus fumigatus delbrlA, delabaA, delwetA, and wild-type RNA-seq |
| SRR5878930 | PRJNA396210 | Aspergillus fumigatus delbrlA, delabaA, delwetA, and wild-type RNA-seq |
| SRR5878941 | PRJNA396210 | Aspergillus fumigatus delbrlA, delabaA, delwetA, and wild-type RNA-seq |
| ERR161596 | PRJEB3185 | Transcription profiling by high throughput sequencing of the pathogenic fungus Aspergillus fumigatus to reveal novel insights in genome structure and MpkA-dependent gene expression |
| ERR161599 | PRJEB3185 | Transcription profiling by high throughput sequencing of the pathogenic fungus Aspergillus fumigatus to reveal novel insights in genome structure and MpkA-dependent gene expression |
| SRR5249268 | PRJNA374516 | Genome-wide transcriptional changes under oxidative stress in iron-starved Aspergillus fumigatus cultures |
| SRR5249270 | PRJNA374516 | Genome-wide transcriptional changes under oxidative stress in iron-starved Aspergillus fumigatus cultures |
| SRR5249269 | PRJNA374516 | Genome-wide transcriptional changes under oxidative stress in iron-starved Aspergillus fumigatus cultures |
| SRR5878939 | PRJNA396210 | Aspergillus fumigatus delbrlA, delabaA, delwetA, and wild-type RNA-seq |
| DRR057055 | PRJDB3180 | Aspergillus fumigatus transcriptome analysis of the asexual developmental stage in atfA deletion mutant |
| DRR057057 | PRJDB3180 | Aspergillus fumigatus transcriptome analysis of the asexual developmental stage in atfA deletion mutant |
| SRR6163648 | PRJNA399754 | In vitro infection of A549 cells with Aspergillus fumigatus |
| SRR6163649 | PRJNA399754 | In vitro infection of A549 cells with Aspergillus fumigatus |
| SRR6163654 | PRJNA399754 | In vitro infection of A549 cells with Aspergillus fumigatus |
| DRR101048 | PRJDB6203 | Transcriptome in response to heat, superoxide, and osmotic stresses in Aspergillus fumigatus mycelia |
| SRR6163669 | PRJNA399754 | In vitro infection of A549 cells with Aspergillus fumigatus |
| SRR5691093 | PRJNA390719 | Characterization of regulators of G protein signaling, rgsD and rax1 of Aspergillus fumigatus |
| SRR5691092 | PRJNA390719 | Characterization of regulators of G protein signaling, rgsD and rax1 of Aspergillus fumigatus |
| SRR5691091 | PRJNA390719 | Characterization of regulators of G protein signaling, rgsD and rax1 of Aspergillus fumigatus |
| SRR5691090 | PRJNA390719 | Characterization of regulators of G protein signaling, rgsD and rax1 of Aspergillus fumigatus |
| DRR075875 | PRJDB5273 | Functional analysis of Aspergillus fumigatus AtrR transcription factor |
| SRR5691088 | PRJNA390719 | Characterization of regulators of G protein signaling, rgsD and rax1 of Aspergillus fumigatus |
| SRR5691089 | PRJNA390719 | Characterization of regulators of G protein signaling, rgsD and rax1 of Aspergillus fumigatus |
| DRR101030 | PRJDB6203 | Transcriptome in response to heat, superoxide, and osmotic stresses in Aspergillus fumigatus mycelia |
| DRR075876 | PRJDB5273 | Functional analysis of Aspergillus fumigatus AtrR transcription factor |
| DRR075872 | PRJDB5273 | Functional analysis of Aspergillus fumigatus AtrR transcription factor |
| SRR5249271 | PRJNA374516 | Genome-wide transcriptional changes under oxidative stress in iron-starved Aspergillus fumigatus cultures |
| SRR5249273 | PRJNA374516 | Genome-wide transcriptional changes under oxidative stress in iron-starved Aspergillus fumigatus cultures |
| SRR5249272 | PRJNA374516 | Genome-wide transcriptional changes under oxidative stress in iron-starved Aspergillus fumigatus cultures |
| SRR5249266 | PRJNA374516 | Genome-wide transcriptional changes under oxidative stress in iron-starved Aspergillus fumigatus cultures |
| SRR5249267 | PRJNA374516 | Genome-wide transcriptional changes under oxidative stress in iron-starved Aspergillus fumigatus cultures |
| SRR5249265 | PRJNA374516 | Genome-wide transcriptional changes under oxidative stress in iron-starved Aspergillus fumigatus cultures |
| DRR075873 | PRJDB5273 | Functional analysis of Aspergillus fumigatus AtrR transcription factor |
| DRR075868 | PRJDB5273 | Functional analysis of Aspergillus fumigatus AtrR transcription factor |
| DRR075869 | PRJDB5273 | Functional analysis of Aspergillus fumigatus AtrR transcription factor |
| SRR5878936 | PRJNA396210 | Aspergillus fumigatus delbrlA, delabaA, delwetA, and wild-type RNA-seq |
| SRR5878933 | PRJNA396210 | Aspergillus fumigatus delbrlA, delabaA, delwetA, and wild-type RNA-seq |
| SRR5878945 | PRJNA396210 | Aspergillus fumigatus delbrlA, delabaA, delwetA, and wild-type RNA-seq |
| SRR5878944 | PRJNA396210 | Aspergillus fumigatus delbrlA, delabaA, delwetA, and wild-type RNA-seq |
| SRR5878935 | PRJNA396210 | Aspergillus fumigatus delbrlA, delabaA, delwetA, and wild-type RNA-seq |
| SRR5878934 | PRJNA396210 | Aspergillus fumigatus delbrlA, delabaA, delwetA, and wild-type RNA-seq |
| SRR5306457 | PRJNA376829 | Transcriptome analysis of Aspergillus fumigatus in the presence of sugarcane bagasse |
| SRR5306455 | PRJNA376829 | Transcriptome analysis of Aspergillus fumigatus in the presence of sugarcane bagasse |
| SRR5306456 | PRJNA376829 | Transcriptome analysis of Aspergillus fumigatus in the presence of sugarcane bagasse |
| DRR075874 | PRJDB5273 | Functional analysis of Aspergillus fumigatus AtrR transcription factor |
| DRR075871 | PRJDB5273 | Functional analysis of Aspergillus fumigatus AtrR transcription factor |
| SRR5438126 | PRJNA382627 | Aspergillus fumigatus strain:CZB01 Transcriptome or Gene expression |
| SRR5439718 | PRJNA382736 | Aspergillus fumigatus Transcriptomes of A1160 and macA deletion strains |
| ERR161597 | PRJEB3185 | Transcription profiling by high throughput sequencing of the pathogenic fungus Aspergillus fumigatus to reveal novel insights in genome structure and MpkA-dependent gene expression |
| DRR075870 | PRJDB5273 | Functional analysis of Aspergillus fumigatus AtrR transcription factor |
| SRR4000584 | PRJNA336568 | Aspergillus fumigatus Velvet temperature response |
| SRR4000585 | PRJNA336568 | Aspergillus fumigatus Velvet temperature response |
| SRR5306452 | PRJNA376829 | Transcriptome analysis of Aspergillus fumigatus in the presence of sugarcane bagasse |
| SRR5306454 | PRJNA376829 | Transcriptome analysis of Aspergillus fumigatus in the presence of sugarcane bagasse |
| SRR5306453 | PRJNA376829 | Transcriptome analysis of Aspergillus fumigatus in the presence of sugarcane bagasse |
| SRR4000590 | PRJNA336568 | Aspergillus fumigatus Velvet temperature response |
| SRR4000591 | PRJNA336568 | Aspergillus fumigatus Velvet temperature response |
| ERR237009 | PRJEB1586 | RNA-Seq of Aspergillus fumigatus mutants resistant to itraconazole |
| ERR237012 | PRJEB1586 | RNA-Seq of Aspergillus fumigatus mutants resistant to itraconazole |
| ERR237007 | PRJEB1586 | RNA-Seq of Aspergillus fumigatus mutants resistant to itraconazole |
| SRR4000588 | PRJNA336568 | Aspergillus fumigatus Velvet temperature response |
| SRR4000589 | PRJNA336568 | Aspergillus fumigatus Velvet temperature response |

Supplementary Figure S3 – Heat maps and clusters of the expression of genes encoding surfome proteins in several different experimental conditions. *A. fumigatus* regulatory networks are publicly available at the National Center for Biotechnology Information (NCBI) and the clustering online tool FungiExpressZ (https://cparsania.shinyapps.io/FungiExpresZ/).
