## Supplementary Figure S4 for "A phylogenetic approach to explore the *Aspergillus fumigatus* conidial surface-associated proteome and its role in pathogenesis"

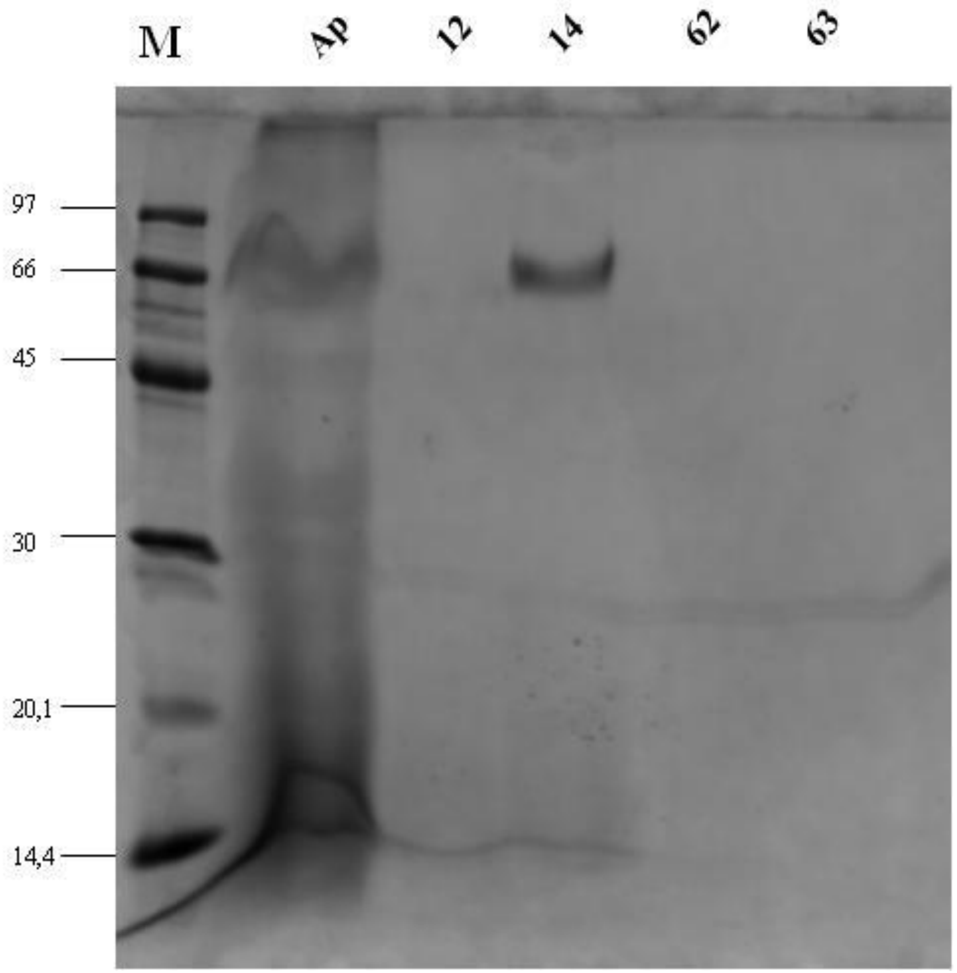


Supplementary Figure S4. SDS polyacrilamide gel showing the purified heterologously expressed AspA in *Pichia pastoris.* Ap, raw extract; fractions 12, 14, 62, and 63.
