## Supplementary Figure S5 for "A phylogenetic approach to explore the *Aspergillus fumigatus* conidial surface-associated proteome and its role in pathogenesis"

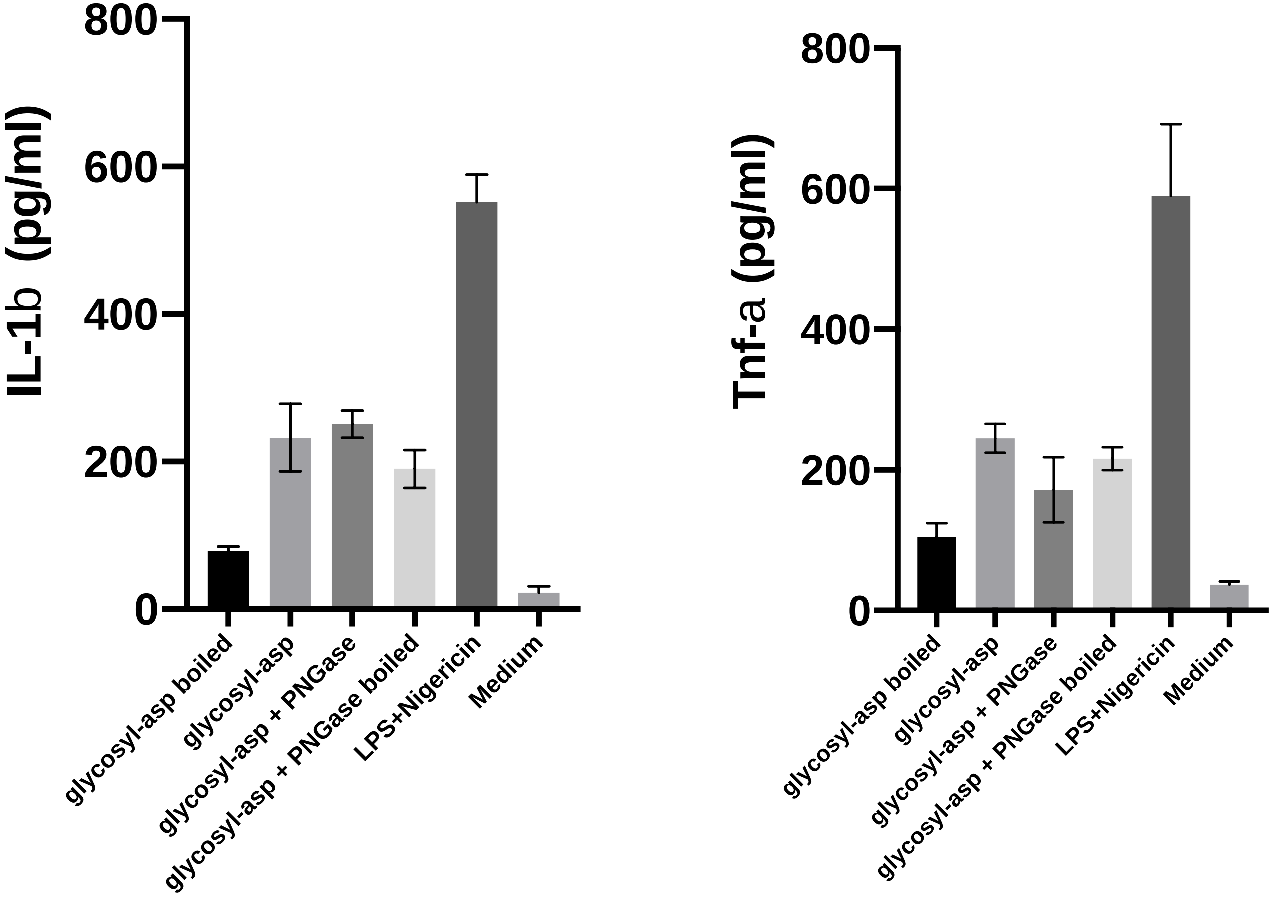


Supplementary Figure S5. IL-1β and TNF-α production in the presence of AspA treated or not with Peptide:N-glycosidase F (PNGase F).
